# Landscape of stimulation-responsive chromatin across diverse human immune cells

**DOI:** 10.1101/409722

**Authors:** Diego Calderon, Michelle L. T. Nguyen, Anja Mezger, Arwa Kathiria, Vinh Nguyen, Ninnia Lescano, Beijing Wu, John Trombetta, Jessica V. Ribado, David A. Knowles, Ziyue Gao, Audrey V. Parent, Trevor D. Burt, Mark S. Anderson, Lindsey A. Criswell, William J. Greenleaf, Alexander Marson, Jonathan K. Pritchard

## Abstract

The immune system is controlled by a balanced interplay among specialized cell types transitioning between resting and stimulated states. Despite its importance, the regulatory landscape of this system has not yet been fully characterized. To address this gap, we collected ATAC-seq and RNA-seq data under resting and stimulated conditions for 25 immune cell types from peripheral blood of four healthy individuals, and seven cell types from three fetal thymus samples. We found that stimulation caused widespread chromatin remodeling, including a large class of response elements shared between stimulated B and T cells. Furthermore, several autoimmune traits showed significant heritability in stimulation-responsive elements from distinct cell types, highlighting the critical importance of these cell states in autoimmunity. Use of allele-specific read-mapping identified thousands of variants that alter chromatin accessibility in particular conditions. Notably, variants associated with changes in stimulation-specific chromatin accessibility were not enriched for associations with gene expression regulation in whole blood – a tissue commonly used in eQTL studies. Thus, large-scale maps of variants associated with gene regulation lack a condition important for understanding autoimmunity. As a proof-of-principle we identified variant rs6927172, which links stimulated T cell-specific chromatin dysregulation in the *TNFAIP3* locus to ulcerative colitis and rheumatoid arthritis. Overall, our results provide a broad resource of chromatin landscape dynamics and highlight the need for large-scale characterization of effects of genetic variation in stimulated cells.

## Introduction

Immune cells respond to extracellular stimuli with stereotyped transcriptional programs that enable their specialized functions during an immune response. These programs, essential for immune homeostasis, are coordinated by precise interactions of transcription factors (TFs) that bind genomic sites to influence chromatin landscape and ultimately gene expression. Tight regulation of these programs is required for an effective and appropriate immune response against cancer and infectious disease, and the avoidance of autoimmunity.

Autoimmune diseases arise when autoreactive immune cells such as B and T cells escape into the periphery and attack host tissues and organs (*1*). During an immune attack, CD4^+^ T cells—which are key players in autoimmune pathology—become activated with self-antigen and environmental stimuli to differentiate into various T helper fates, including effector Th1, Th2 and Th17 cells (*2*). These CD4^+^ T cell subsets secrete inflammatory cytokines to elicit distinct responses, resulting in persistent inflammation, cellular infiltration and ultimately tissue damage. Each subset is regulated by distinct *cis-* regulatory enhancer landscapes and transcriptional programs that are controlled by unique sets of TFs. Thus, genetic variation in regulatory regions that tune these transcriptional programs can contribute to the risk of human autoimmune disease.

Genome-wide association studies (GWAS) have identified hundreds of genetic variants that contribute to the risk of autoimmunity. Roughly 90% of these signals lie in non-coding regions, and thus presumably act by altering gene regulation, however most of these remain difficult to interpret. A number of studies have illustrated enrichment of variants linked to risk of immune-mediated disorders at key enhancers and cell-specific expression quantitative trait loci (eQTLs), suggesting potential mechanisms by which non-coding variants contribute to disease pathology (*3-9*). Nonetheless, only a minority, perhaps 25% (*10*), of GWAS signals can currently be explained through known eQTLs.

Several groups have shown that additional GWAS-eQTL overlap can remain hidden within stimulation-specific functional regions of immune cells (*4, 11-15*). Probing these response-specific functional regions can reveal previously undetected disease-associated mechanisms, emphasizing the unique role of stimulation to autoimmunity. For example, our group recently discovered a novel stimulation-response regulatory element which, when perturbed, resulted in a dysregulated immune response by delaying IL2RA expression *in vivo* (*16*). Furthermore, this region harbors a fine-mapped GWAS variant linked to inflammatory bowel disease and type 1 diabetes (*17-19*), thus connecting immune response to autoimmunity. However, these studies were performed on relatively few stimulated cell types. At present we lack a comprehensive view of the effects of stimulation on the chromatin landscape of diverse immune cell types, and the role of SNPs in these regions on autoimmune disease.

To address this gap, we developed a comprehensive resource of chromatin accessibility and gene expression from 25 primary human immune cell types isolated from four human blood donors, in both resting and activated states. In addition, to further understand chromatin structure in cells important for T cell development we included six thymocyte subsets and thymic epithelial cells (TECs) collected from three fetal thymus samples.

Overall, we observed features of the chromatin landscape that depend upon cell lineage, and also large-scale responses to stimulation. Notably, B and T cell subsets shared a significant proportion of these stimulation-responsive chromatin regions. Integrating these data with autoimmune GWAS, we found that stimulation-responsive chromatin regions explained significant trait heritability in multiple immune cell types, indicating distinct lineage contributions to autoimmunity. Finally, we leveraged our newly characterized stimulation-responsive chromatin elements and allele-specific imbalance of chromatin accessibility as a functional readout of variants from individual donors to identify new autoimmune-related mechanisms. Using this approach, we fine-mapped a SNP that is associated with both rheumatoid arthritis and ulcerative colitis (rs6927172), as a potential causal variant that likely regulates the expression of *TNFAIP3*. Taken together, our study shows the importance of broadly shared stimulation chromatin regions in the context of autoimmune disorders and provides a resource for understanding unique cell type regulation networks.

## Results

### A functional atlas of diverse human immune cell types in resting and stimulated states

To identify epigenomic elements underlying cellular differentiation and stimulation responses in the human immune system, we generated a map of genome-wide chromatin accessibility and gene expression of the human immune system in resting and stimulated cells (Figure 1A). We isolated 25 distinct, specialized immune cell types by flow cytometry from peripheral blood of up to four healthy donors including different subsets of B cells, CD4^+^ T cells, CD8^+^ T cells, γδ T cells, monocytes, dendritic cells (DCs) and natural killer cells (NK) (Figure S1, Table S1). Additionally, we collected fetal thymus samples from three donors and isolated a number of thymocyte subsets, including double positive (DP), double negative (DN), pre-T double negative (pre-T DN), immature single positive CD4^+^ (CD4^+^ iSP) and single positive mature CD4^+^ and CD8^+^ T cells, as well as thymic epithelial cells (TEC).

**Figure 1.**
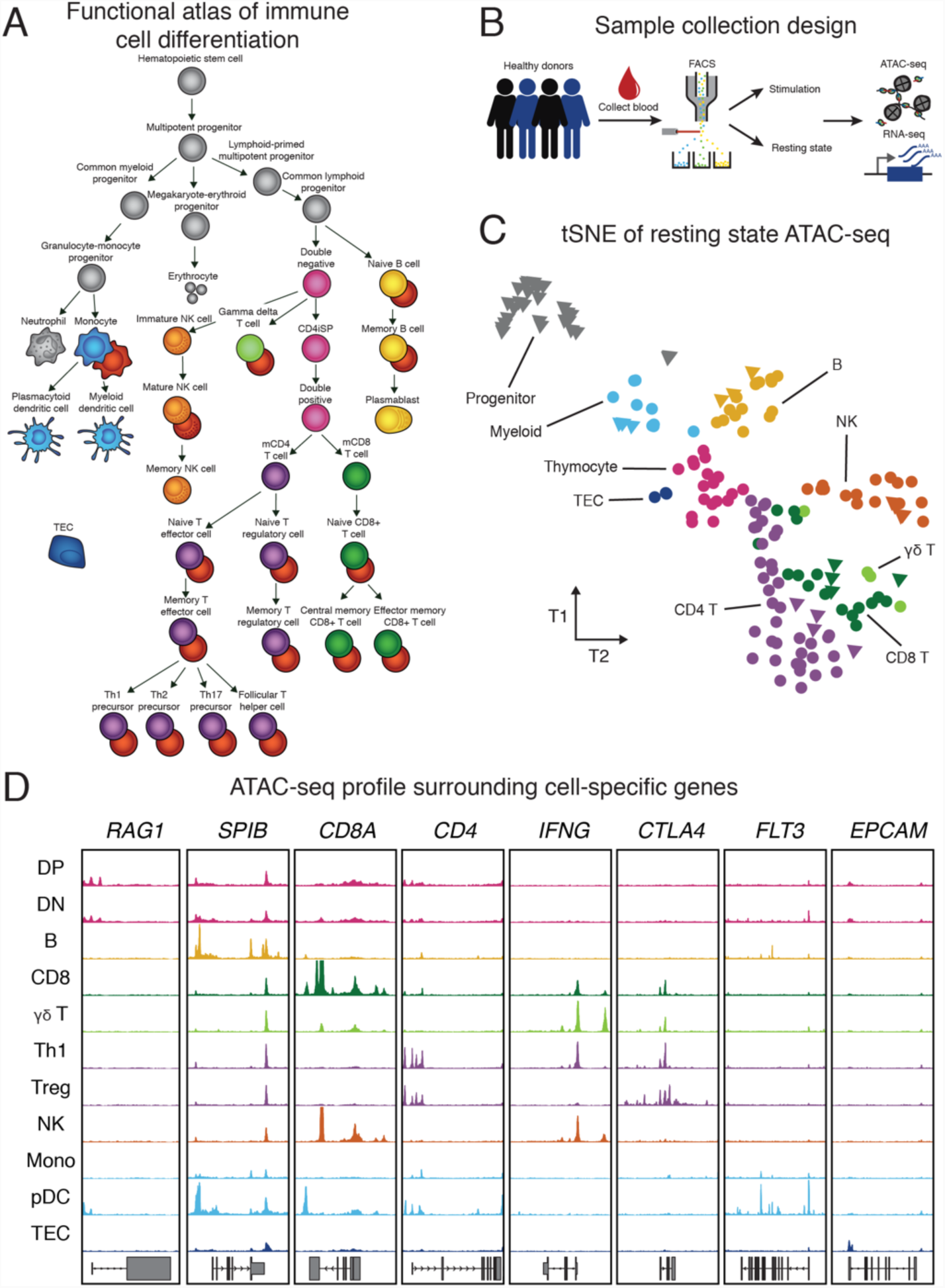
Study workflow and tSNE of ATAC-seq data. (A) Illustration of isolated cell types, which include cell types that were previously published (gray), and resting (colored) and stimulated (red) immune cells obtained from this study. (B) Schematic of sample processing pipeline. Immune cells were sorted by flow cytometry from up to four healthy donors, activated and subjected to ATAC-seq and RNA-seq. Not pictured are the fetal thymocytes and thymic epithelial cells we collected. (C) Exploratory tSNE on ATAC-seq chromatin accessibility of all cell types under resting state. Each sample is colored by broad cell lineage. Biological replicates for each cell type (n = 3-4) are plotted separately. Triangles represent previously published data (*20*) and circles represent data generated in this study. (D) Representative ATAC-seq profiles (y-axis scales go from 0 to 400 RPKM) at several cell type-specific genes (see Methods).

We performed an assay for transposase-accessible chromatin using sequencing (ATAC-seq), which profiles chromatin-accessible regions as a sequencing depth read out (*21*), and RNA-seq on all cell types isolated from whole blood. We observed high technical reproducibility (ATAC and RNA mean Pearson’s R value = 0.89 and 0.80, respectively) and biological reproducibility (ATAC and RNA, mean Pearson’s R value = 0.85 and 0.77, respectively) across replicates. We further confirmed the quality of our data by analyzing the enrichment of ATAC-seq reads mapping to transcription start sites (TSS) and the expression of cell type-specific genes. As expected, we observed strong enrichment of reads at TSSs genome-wide and at promoters of cell type-specific genes such as *CD8A* in CD8^+^ T cells, consistent with highest expression of *CD8A* in the corresponding cell type compared to others (Figure 1D, Figure S2A, B). On average, across all resting cell type samples from multiple donors, we identified 36,810 accessible peaks (q < 5%), controlling for the effects of read depth and sample quality.

Next, we characterized global features of chromatin accessibility across resting cell types. Unsupervised clustering of ATAC-seq profiles revealed distinct chromatin signatures for different cell lineages (Figure 1C, Figure S2C). For example, hematopoietic stem cell progenitors clustered separately from more differentiated immune cell types (data from 20). T cell subsets including CD4^+^, CD8^+^ and γδ T cells clustered closely together, indicating similar chromatin accessibility profiles for each of them under resting conditions. Moreover, while circulating mature T cells originate from single positive thymocytes, they could be distinguished from their precursors based on distinct accessibility profiles, suggesting that mature T cells have undergone further chromatin remodeling in the periphery. Finally, where available, the samples generally clustered as expected with published data from the same cell types (Figure 1C). The same major cell type clusters observed in the chromatin accessibility data were recapitulated when we clustered the RNA-seq samples (Figure S2C,D). However, we found that cell type clustering accuracy was higher when using chromatin accessibility than when using gene expression (mean HA-adjusted RAND index 0.19 and 0.12 respectively), consistent with previous analyses (*20*).

### Stimulation results in large-scale chromatin and gene expression changes

Previous studies of individual cell types have shown large-scale chromatin remodeling upon stimulation (*11, 12, 22, 23*). We set out to perform a comprehensive analysis of stimulation-dependent chromatin and gene expression changes across a wide range of cell types and lineages. To investigate these effects, we *ex vivo* stimulated the majority of the collected cell types and performed ATAC-seq and RNA-seq.

Overall, we found that stimulation drives dramatic changes in regulatory landscapes of B and T cells. In contrast, we saw limited effects in innate lineage cells (Figure S3A,B), perhaps because the chromatin response in the innate lineage cells is poised or transient (*22, 24, 25*).

Stimulation was a major driver of sample clustering of B and T cells (Figure 2A). The primary axes of variation in chromatin accessibility across cell types of the adaptive lineage were associated with both stimulation and cell type. Moreover, stimulated samples moved in a similar direction, suggesting an underlying shared chromatin response to stimulation.

**Figure 2.**
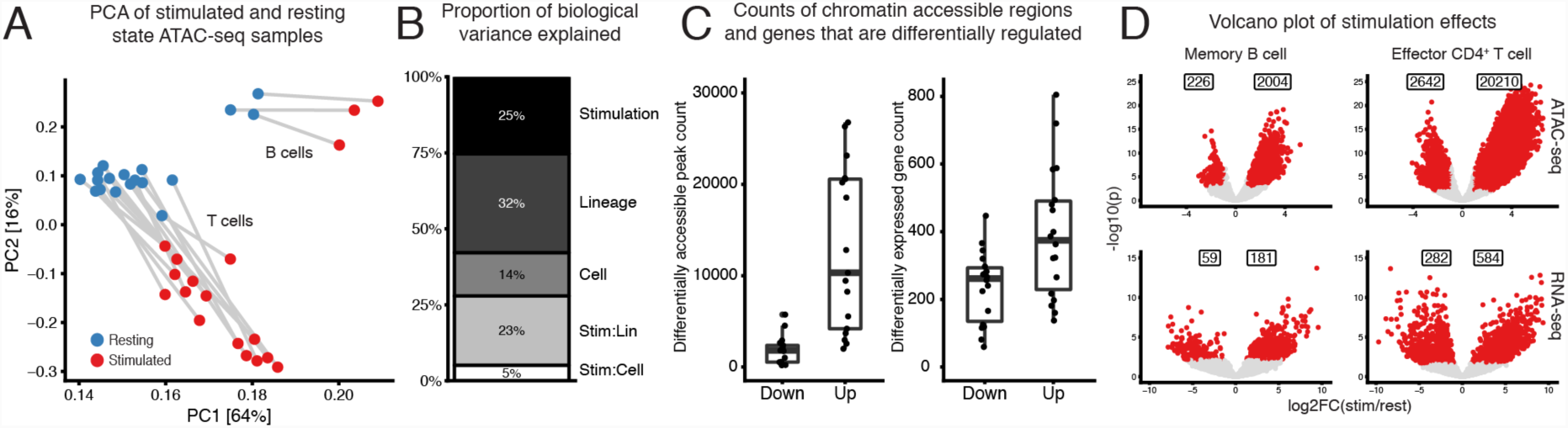
Stimulation induces large-scale changes in chromatin and gene expression in B and T cells. (A) PCA of ATAC-seq read counts of B and T cell subsets (excluding plasmablast cells), which were merged from multiple donors (n = 3-4), including resting (blue) and stimulated samples (red). Analysis based on the top 100k most variable peaks. (B) We used a random effects model to estimate the proportion of variation in chromatin accessibility, of the samples in (A) with at least 3 biological replicates, explained by biological factors of interest (see Methods). The interaction effect for lineage is labeled Stim:Lin and the interaction effect for cell type is labeled Stim:Cell. Biological effects explained 25% of the chromatin variation.(C) Counts of significant differentially accessible chromatin regions (left panel) and expressed genes (right panel) identified for B and T cells during stimulation. Naïve regulatory T cells were excluded because we had too few biological samples passing QC to identify many significant accessibility regions. (D) Example of volcano plots showing stimulation effect estimates for ATAC-seq (top) and RNA-seq (bottom) for memory B (left) and effector CD4+ T (right) cells. Significant (absolute log2FC > 1, q < 1%) differentially accessible regions and expressed genes are indicated in red and the counts shown in the boxes above.

To quantify these observations, we used a random effects model to estimate the proportion of biological variance of chromatin accessibility explained by stimulation condition, lineage (CD4^+^, CD8^+^, and B) and cell subset (e.g. naïve, memory and T helper cells)(see Methods). We found that stimulation, regardless of lineage or cell type, accounted for a quarter of the explained chromatin variation (Figure 2B). In fact, the chromatin differences observed due to stimulation were nearly as large as differences between cell lineages (25% versus 32%, respectively). Additionally, lineage-dependent stimulation changes explained a significantly greater amount of chromatin variation than cell subset-specific stimulation. In summary, broadly shared simulation and lineage-specific stimulation effects drove a large proportion of observed chromatin variation.

One major effect of stimulation was a marked increase in the number of accessible sites and in widespread up-regulation of gene expression (Figure 2C). Specifically, as identified by MACS2 (q < 5%) and correcting for confounding effects (see Methods), we found 30,224 additional peaks in stimulated cell types compared to resting state (Figure S3C). Testing for differential accessibility, we found, on average, that 12,119 (9,878 median) peaks were significantly more accessible, versus just 1,722 (1,781 median) peaks that were significantly less accessible following stimulation (Figure 2C, left panel) (Table S2). We observed a similar pattern in gene expression where, per sample, an average of 394 (median 375) genes were up-regulated compared to 237 (median 262) down-regulated genes in stimulated cell types relative to their resting cell state (Figure 2C, right panel and Table S3).

For example, stimulation of memory B cells leads to increased accessibility for 2,004 peaks, and increased expression of 181 genes (Figure 2D). At the same time, only 226 peaks become less accessible and 59 genes are down-regulated following stimulation. Similar to B cells, we found the same trend within the T cell lineage: for example, effector CD4^+^ T cells have 20,210 peaks with increased accessibility following stimulation compared to only 2,642 peaks with decreased accessibility. Overall, these results illustrate the global chromatin and transcriptional changes of immune cells upon stimulation. In addition to the chromatin state of a cell under resting conditions, new stimulation networks become activated, resulting in increased chromatin accessibility and gene expression.

We next sought to identify transcription factors that may drive cell type and stimulation-responsive elements. We investigated variation in accessibility at position weight matrix (PWM)-predicted TF binding regions across cell types and conditions (Figure S3D). For example, the PU.1 motif is most enriched in B cells, DCs, and monocytes, consistent with gene expression data (Figure S3E). The corresponding TF for the PU.1 motif is integral to both myeloid and lymphoid B cell development (*26*).

In contrast, we found that the BATF motif was consistently enriched in accessibility regions across stimulated samples compared to their corresponding unstimulated state (Figure S3D). This suggests a shared effect of BATF and/or related transcription factors on chromatin regulation in stimulated samples across cell lineages. Additionally, the putative activity of BATF correlates with upregulation of *BATF* expression (Figure S3F) and the expression of several classes of previously identified BATF-target genes (Figure S3G) (*27*). Thus, our analysis identified large-scale genome-wide changes in chromatin accessibility and gene expression upon stimulation in B and T cells putatively attributable to specific TFs.

### Stimulation responsive elements are broadly shared across B and T cell lineages

Given that both B and T cells have highly distinct functions during an immune response, we were interested in determining whether stimulation of different lineages gives rise to unique or shared components of chromatin remodeling. Therefore, we next sought to estimate the prevalence of stimulation effects that were shared or unique between any two cell types.

For example, we observed regions linked to stimulation response that were uniquely open in either B or CD8^+^ T cells but not both (Figure 3A, left panel). At the *BACH2* locus, which encodes a key factor in driving T cell differentiation (*28*), we observed a site upstream of the gene promoter that only becomes accessible in CD8^+^ T cells upon stimulation. Additionally, we detected stimulation-accessible regions that were shared between B and CD8^+^ T cells, including *JMJD1C* (Figure 3A, right panel). Generalizing these observations genome-wide, we found that estimates of stimulation effects were correlated between CD8^+^ and effector CD4^+^ T cells and even CD8^+^ and B cells (Figure 3B). Thus, to quantify the prevalence of a shared stimulation response we computed the correlation of effects estimated from stimulation-induced changes in chromatin accessibility and gene expression between each pair of cell types (Figure 3C).

**Figure 3.**
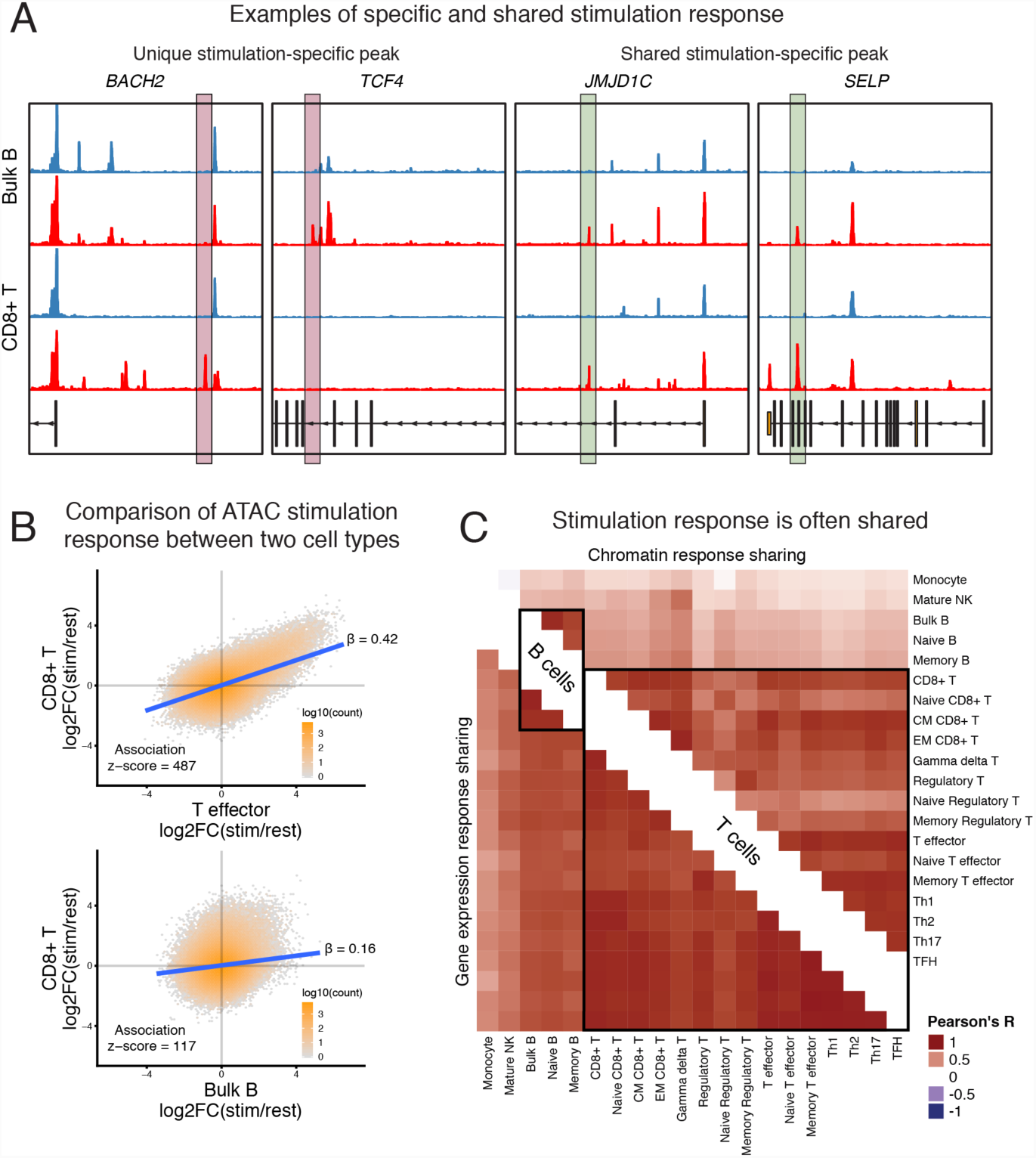
Quantifying the shared stimulation response across cell lineages. (A) Representative ATAC-seq profiles (y-axis scale is from 0 to 400 RPKM) of unique (left) and shared (right) stimulation-specific chromatin regions for bulk B (top) and CD8^+^ T cells (bottom). Resting and stimulation states are colored in blue and red, respectively. The peaks of interest are highlighted. (B) Example scatter plots showing comparisons of log2 fold change estimates of the response of chromatin accessibility to stimulation. High-density regions are indicated in orange. (C) Quantification of the shared stimulation response. Displayed is the Pearson’s R correlation between samples from stimulation-response chromatin (top-right triangle) and gene expression (bottom-left triangle) effects, at sites or genes with a significant stimulation response in at least one of the two cell types in the comparison. All estimates are from at least three biological replicates, except the Naïve Regulatory T cells, which had two.

We observed large proportions of shared stimulation-induced chromatin accessible regions between CD4^+^, CD8^+^ and γδ T cells (mean R = 0.74) upon activation. Similarly, we observed strong sharing among B cell subsets (mean R = 0.82). Together, these observations suggest that CD3/CD28 signaling, which are responsible for stimulation response in T cells, and B cell receptor and IL-4 signaling which drive stimulation response in B cells, both induce dramatic and consistent patterns of chromatin remodeling within B and T lineages, respectively.

Furthermore, despite the divergent mechanisms of activation in B and T cells, we identified a significant level of sharing globally between stimulation-response in B and T cells (mean R = 0.37), indicating that some stimulation-induced regulatory networks are utilized by both B and T cells. Finally, while stimulation induces both unique and shared chromatin responses, the stimulation-induced transcriptional signatures are broadly shared among cell types (mean R = 0.77) (Figure 3D). Thus, we have identified a strong, shared component of stimulation response that is broadly active in various T and B cell subsets.

### Allelic imbalance as a tool to study context-specific regulation

In addition to identifying stimulation-associated chromatin regions, we were interested in characterizing genetic variants that alter context-specific chromatin regulation. Due to the small number of individuals included in this study, and thus insufficient power to call chromatin accessibility quantitative trait loci (caQTLs), we used the observed allelic imbalance of ATAC-seq reads that map to heterozygous SNPs to identify significant sites of allele-specific chromatin accessibility (ASC). The hypothesis is that a heterozygous variant may cause local, allele-specific changes in chromatin accessibility, for example by disrupting local transcription factor binding. Such events result in an imbalance in the number of ATAC-seq reads overlapping each allele (*29-33*). After read-filtering to account for mapping biases (*34*), we identified 607 significant ASC sites, on average, per sample, and a total of 10,780 sites overall (q < 10%, see Methods for details, Table S4)

Because we collected many cell samples from each donor, we could test for allelic imbalance of chromatin accessibility at an individual heterozygous site across cell type and condition-specific contexts. Thus, our data enabled us to identify sample- and context-specific ASC sites, respectively. For instance, we observed a heterozygous mutation, rs3795671 that resulted in increased chromatin accessibility on the reference allele across resting and stimulated cell types from all collected lineages (Figure 4A). In contrast, other variants only displayed measurable allelic imbalance in a subset of cell type states; these include rs7011799, rs12091715, rs1250567, and rs1445033, which are associated with chromatin accessibility within resting, stimulated, T, and B lineage contexts, respectively.

**Figure 4.**
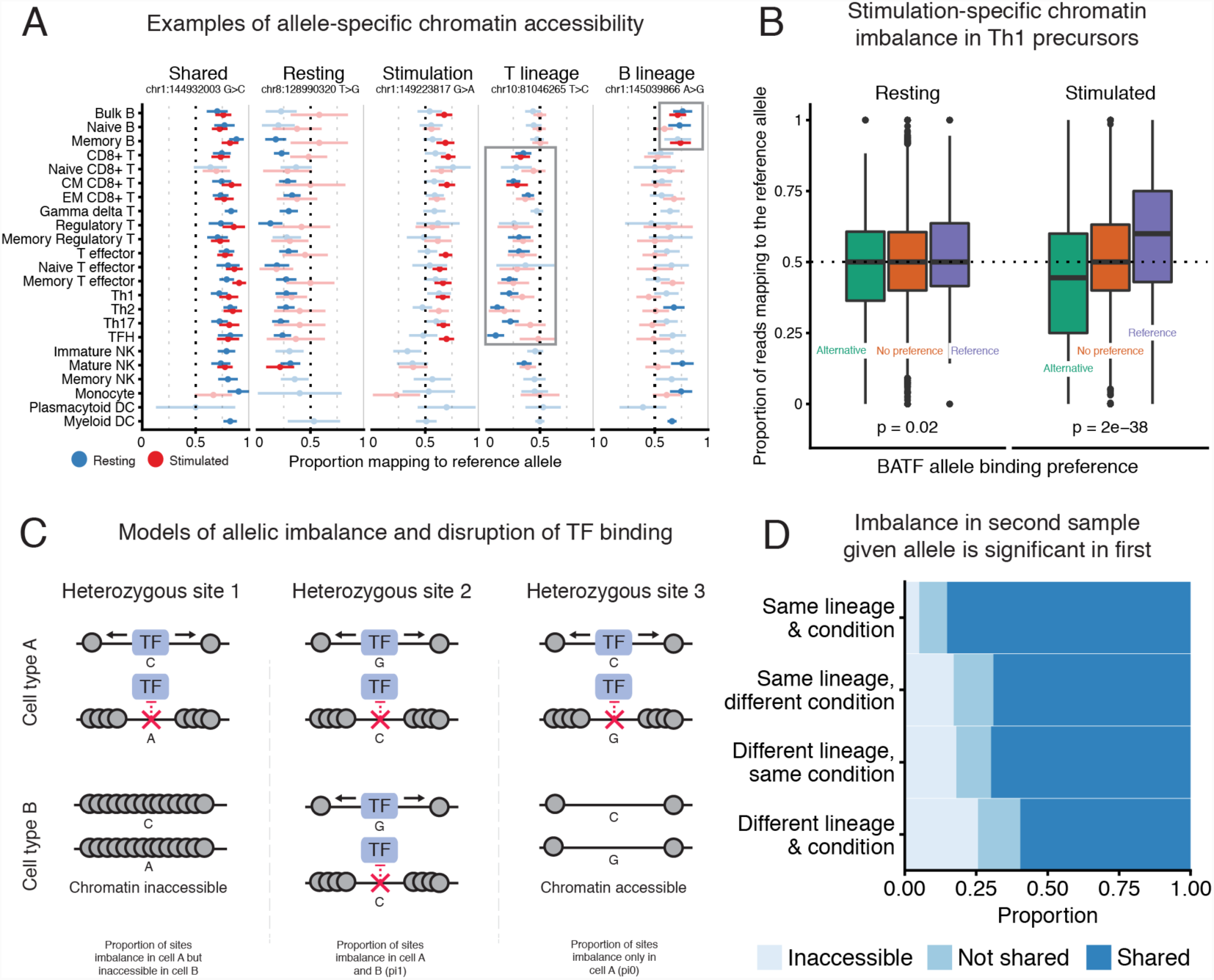
Observed allelic imbalance in chromatin accessibility data. (A) Examples of allele-specific chromatin accessibility imbalance shared across various groups. For each heterozygous site, we display the proportion of reads mapping to the reference allele (x-axis) for resting (blue) and stimulated (red) cell samples (y-axis). Error bars represent 95% confidence intervals and were computed from read depth. Samples without significant imbalance are lightly shaded. We excluded samples with fewer than four reads. (B) We grouped all heterozygous sites into three groups based on the PWM-predicted BATF binding affinity: preference for the alternative (green), no preference (orange), preference for the reference allele (purple). Next, we displayed the aggregate proportion of reads mapping to the reference allele for these groups in Th1 precursor cells under resting (left) and stimulated (right) conditions. We excluded sites with fewer than four reads. (C) Model for possible scenarios of allele-specific chromatin accessibility imbalance in two different cell types or conditions for the same donor. In all cases, we condition on the fact that there is a heterozygous variant that disrupts the binding of a chromatin-regulating factor in cell type A (top). In cell type B, we can observe one of three possibilities, from left to right: 1) chromatin is inaccessible, 2) the heterozygous site disrupts chromatin regulation (referred to as shared effect), or 3) the chromatin is accessible, but there is no chromatin imbalance detected. (D) Proportion plot displaying the estimated average proportions for each case from (C), stratified by whether the two samples were of the same lineage and condition. We excluded innate cells from this analysis. All plots are for Donor 1, who had the highest sequencing depth.

Next, we were interested in leveraging these data to verify TFs that regulate chromatin in specific contexts, and thus drive context-specific allelic imbalance. As described above, motif enrichment analysis (Figure S3D) suggested that BATF, among others, is an important regulator of chromatin in stimulated cells (*23*). We therefore hypothesized that sites that affect BATF binding would result in ASC within stimulated samples. As a proxy for true BATF binding, for each heterozygous site within a BATF binding motif, we computed the relative binding affinity for the reference and the alternative allele using the BATF PWM. Upon stimulation, alleles that were predicted to increase BATF binding affinity were associated with increased accessibility (ASC>0.5). Strikingly however, there was no such effect in resting cells, indicating that BATF does not impact chromatin accessibility in the resting state (Figure 4B). We observed a similar preferential binding effect for other stimulation-associated TFs, including JUNB, FOSL1 and BACH1 (Figure S4). Thus, the PWMs of these factors can predict sequence dependent changes of stimulation-specific effects on chromatin accessibility.

Intrigued by the stimulation-specific effect of BATF, we wanted to leverage our ASCs to determine the prevalence of context-specific chromatin regulation. We propose three possible models of chromatin effects at a significant ASC site, in pairs of cell types or conditions (Figure 4C, from left to right). The variant can: i) differentially affect chromatin in cell type A, while the region is inaccessible in cell type B; ii) differentially affect chromatin in both cell types, or; iii) differentially affect chromatin in cell type A, but not in cell type B, even though the region is accessible therein.

We quantified the proportion of ASCs that fit each possible model and observed that most ASCs have shared effects on chromatin regulation (see Methods). To further investigate changes in the prevalence of shared regulation across differences in cell state, we varied whether the two compared samples were of the same lineage and condition. Notably, most effects on allelic imbalance are shared regardless of lineage or condition (Figure 4D). In contrast, the proportion of accessible sites covaried more strongly with differences in lineage and condition between the two samples. These observations suggest that changes in cellular function due to chromatin remodeling occur primarily through changes in chromatin accessibility, rather than regulation of already accessible chromatin. The BATF analysis in Figure 4B indicates that at least some sites containing BATF motifs are likely differentially regulated upon stimulation, however this scenario represents a small minority of all ASC sites.

### GWAS enrichment in immune-specific chromatin accessible regions

Understanding the molecular mechanisms behind autoimmune disease risk variants requires the identification of disease-relevant cell types and states. With chromatin accessibility information from a large number of resting and activated immune cell types, we can identify cell types and conditions enriched for variants associated with autoimmune disease. Using publicly available summary statistics for nine autoimmune disorders and four non-immune traits (as controls), we identified trait-relevant functional annotations using the partitioning heritability functionality of LD Score regression (*3*). This allowed us to compute the proportion of heritability explained by SNPs in open chromatin for a particular cell type. This quantity, divided by the overall proportion of open chromatin SNPs in that cell type, will be referred to as the enrichment of heritability (Table S5).

For most autoimmune traits, we observed greater enrichment of heritability in differentiated immune cell types compared to progenitor cells and non-immune tissues (Figures 5A and S5A,B). For example, for rheumatoid arthritis (RA) we saw about ten-fold enrichment of heritability in open chromatin from control tissues and immune progenitors compared to the genome-wide background (Figure 5A), and a median 23-fold enrichment in immune cell open chromatin depending on the cell type. The enrichment in non-immune tissues compared to the genome-wide background is likely due to the fact that many open chromatin regions are shared among diverse cell types. Thus, many of the SNPs that affect disease risk through immune cell functions are also located in open chromatin in many other tissues (*35*).

**Figure 5.**
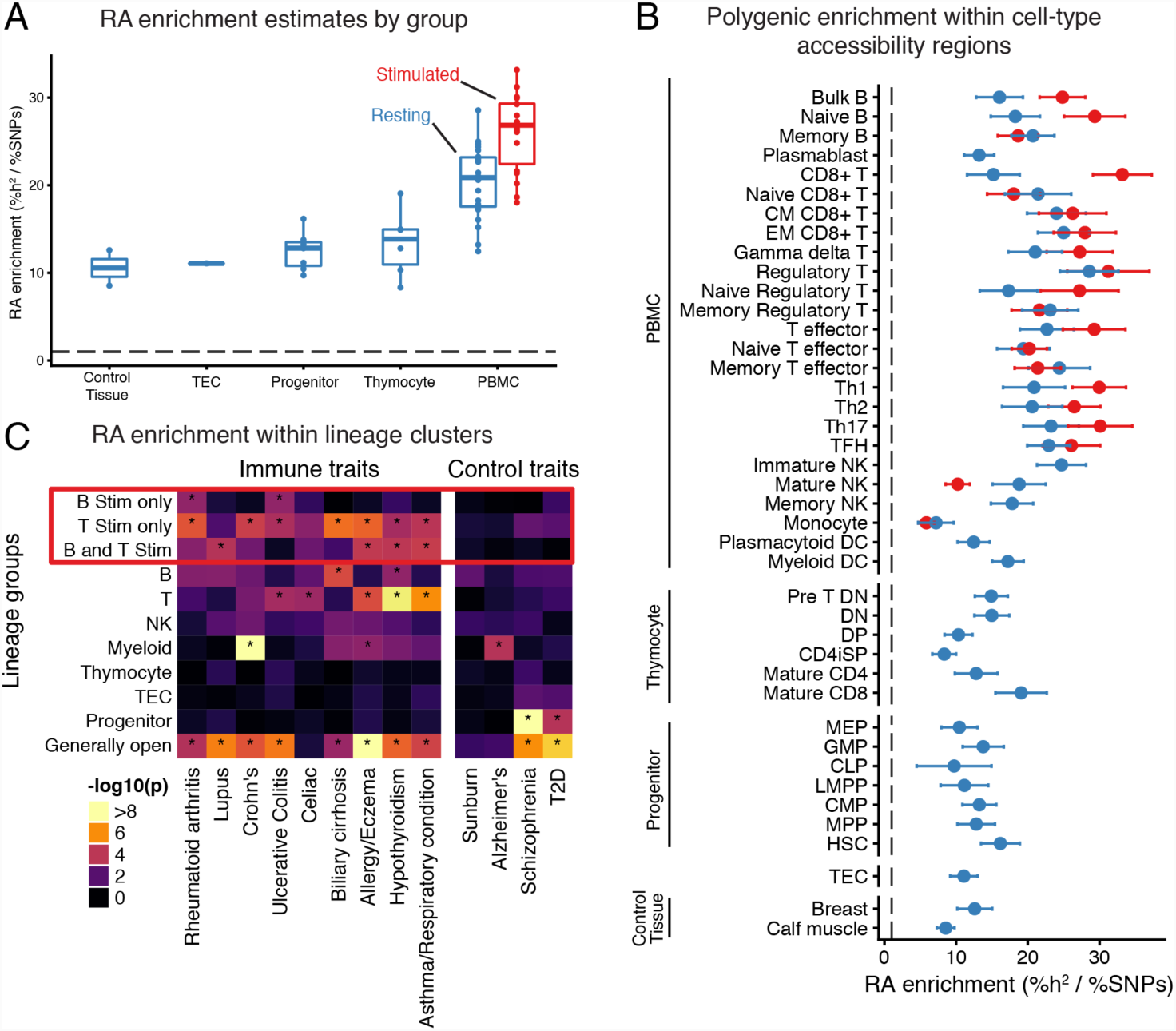
GWAS analysis of accessible regions. (A) Rheumatoid arthritis heritability enrichment is aggregated by differentiation and condition. Stimulated innate cells were excluded from this visualization. (B) Enrichment of rheumatoid arthritis (RA) heritability (x-axis) in open chromatin regions for resting (blue) and stimulated (red) samples (y-axis). Error bars indicate one standard deviation in each direction. (C) We grouped peaks into disjoint clusters based on their patterns of accessibility across cell types (x-axis). Then, we used these peak clusters (y-axis) to identify enrichments of trait signal (y-axis). We highlighted groups of peaks related to stimulation with a red box, and asterisks (*) indicate significant enrichment of trait heritability (Bonferroni adjusted p < 0.05).

Notably, the heritability enrichment was particularly strong in stimulated immune cells, compared to their resting counterparts (Figure 5B). This signal was spread across diverse cell types, including both B and T cell lineages, and did not implicate particular immune subsets as central drivers of RA risk. We wondered whether this relatively diffuse signal arises because multiple cell types play causal roles in RA, or simply because the broad sharing of stimulation peaks makes it difficult to identify a critical cell type (Figure 3C).

To investigate this further, we grouped the open chromatin peaks into 11 disjoint clusters based on their profiles of accessibility across cell types and conditions (see Methods, Figure S5C). For example, we defined a cluster of peaks that are open only in stimulated B cells, and a cluster of peaks that are open only in stimulated T cells. We used these peak groupings to test which clusters are enriched for autoimmune trait heritability (Figure 5C).

Overall, we observed strong enrichment of heritability in accessible regions from multiple peak groups, especially among the stimulation-related peaks. No single peak group could explain the entirety of the signal. Since these groups are disjoint, this observation implies that multiple separate immune components contribute to heritability. In the case of RA, we identified enriched heritability in stimulation peaks specific to B lineage cells, and in stimulation regions specific to T lineage cells, thus implicating both stimulated B and T cells. Interestingly, previous studies have separately identified B (*20, 36*) and T cells (*4, 6*) in driving the development of RA, although our analysis would suggest that both contribute to autoimmunity. In addition to signals from cell type-specific clusters, all traits had significant enrichment within loci that were broadly accessible, highlighting the contribution of broadly open “housekeeping” peaks to heritability.

Taken together, these results suggest that stimulated cells from both B and T lineages likely contribute to autoimmunity. Furthermore, it remains difficult to implicate more narrowly-defined subsets through this type of analysis, due to the extensive sharing of open chromatin among closely related subsets.

### Stimulation and cell type-specific functional data increase GWAS and eQTL overlap

One important approach for linking noncoding GWAS hits to the genes they regulate is through eQTL mapping. However, only a small proportion of significant GWAS sites have been successfully linked to an eQTL. One hypothesis for the limited overlap is that many eQTLs may be context-specific and thus remain unidentified if not all of the relevant cell types or conditions have been studied (*37*). Since we found a strong signal of autoimmune trait risk variants located within stimulation-specific chromatin accessibility peaks, we sought to investigate whether these variants would have been identified as candidate regulatory variants in a standard tissue-based eQTL study. Here we present results primarily from RA, because the heritability of this disorder was highly enriched in the stimulation-specific accessible regions in our previous analysis.

For this analysis, we partitioned the set of allelic imbalance sites into two groups: ASC sites that were identified in resting samples, and ASC sites that were only identified in stimulated samples (either B or T cells; nominal p < 0.10). In this analysis, sites that were present in both resting and stimulated cells were included in the resting set, as those could be identified even without using stimulated cell data. Since variants that affect chromatin accessibility are strongly associated with regulation of gene expression (*29*), the former set of ASCs likely contain sites that regulate gene expression in the resting cell state, while the latter contain sites associated with stimulation-specific gene regulation.

Both sets of SNPs exhibited highly significant enrichment of RA GWAS signal relative to control SNPs (p = 2 × 10^-19^ (resting), and p = 3 × 10^-14^ (stimulation only); Figure 6A). However, importantly, the stimulation-specific ASCs would not have been identified in resting samples. Thus, stimulation-specific chromatin is as important, in terms of disease risk enrichment, as resting-specific chromatin from immune cells

**Figure 6.**
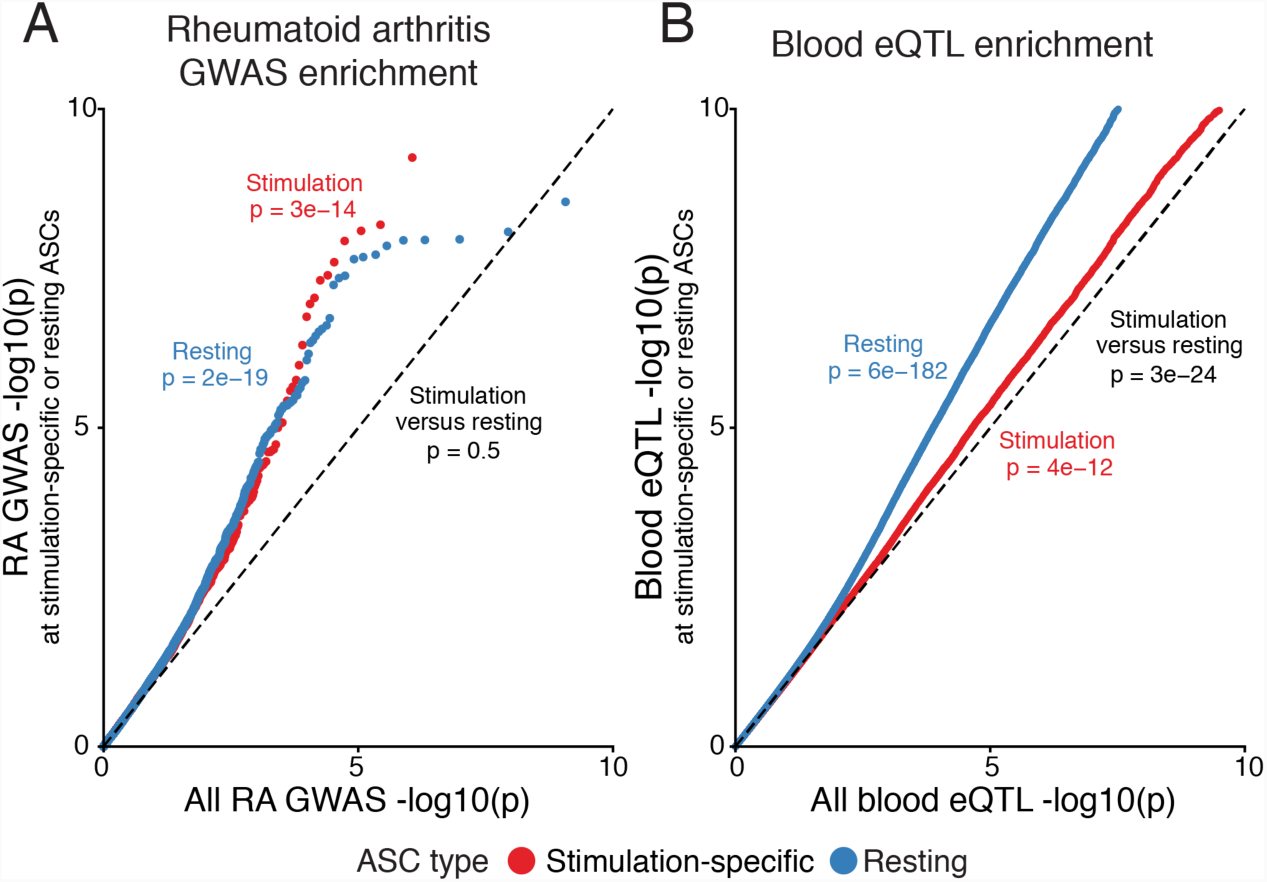
GWAS and eQTL enrichment in sites of allele-specific chromatin. (A) Comparison of rheumatoid arthritis GWAS enrichment within the set of SNPs that regulate chromatin accessibility, in either B or T cells, under stimulation (red) or resting (blue) conditions. (B) Comparison of eQTL signal in the same two sets of variants from (A), using eQTL data from GTEx v7. For both plots the x-axis reflects an empirical distribution of p-values.

Thus far there have been many studies of blood eQTLs (PBMCs) encompassing many thousands of samples (*38*). However, since the proportion of stimulated cells in bulk tissues is relatively low, we hypothesized that sites that affect stimulation-specific chromatin accessibility may not be detected as eQTLs using whole blood or resting immune cells. To test this hypothesis, we determined the enrichment of blood eQTL signal from GTEx in the two previously described sets of resting and stimulation-specific ASC sites. We observed a strong enrichment (p = 3 × 10^-24^) of blood eQTL signal within the resting state-specific set of ASC variants compared to the stimulation-specific set (Figure 6B). These observations indicate a clear need for large-scale eQTL mapping in stimulated immune cells, and argue that the low overlap of autoimmune GWAS and eQTL data (*10*) is at least partly driven by the current lack of such data.

Together these observations suggest that the inclusion of such data from stimulated cells could help fine-map previously unexplained causal mutations. To scan for such examples, we intersected a database of fine-mapped candidate causal GWAS SNPs (*4*) with a set of stimulation-specific allelic imbalance heterozygous SNPs and PWM-disruption scores for TFs (see Methods). This approach identified a T cell lineage-specific stimulation peak within an intergenic region upstream of the *TNFAIP3* locus (encoding A20 protein). Moreover, this region contains several variants that are included in a credible set of regions identified for RA and ulcerative colitis (UC). For RA, only rs6927172 falls within a stimulation-specific chromatin accessible peak (Figure 7A). For UC, in addition to rs6927172, three additional variants fall within another nearby stimulation-responsive peak, yet this second peak does not demonstrate as strong of a response as the first peak (Figure S6A). Notably, we found that rs6927172 was significantly associated with altered chromatin accessibility in stimulated CD4^+^ T cells across multiple donors, suggesting a potential role of the SNP in regulating chromatin remodeling (Figure 7B). Indeed, the cytosine to guanine mutation leads to a large change in the PWM-predicted binding affinity of the p^50^ subunit of NF?B1 (Figure 7C, Figure S6B; see FigureS6C for expression levels). As NF?B1 has been shown to regulate *TNFAIP3* expression (*39*), this result suggests that rs6927172 may contribute to the pathology of autoimmune diseases by disrupting binding of NF?B1 in stimulated T cells to the *TNFAIP3* locus and therefore inhibit gene expression (Figure 7D).

**Figure 7.**
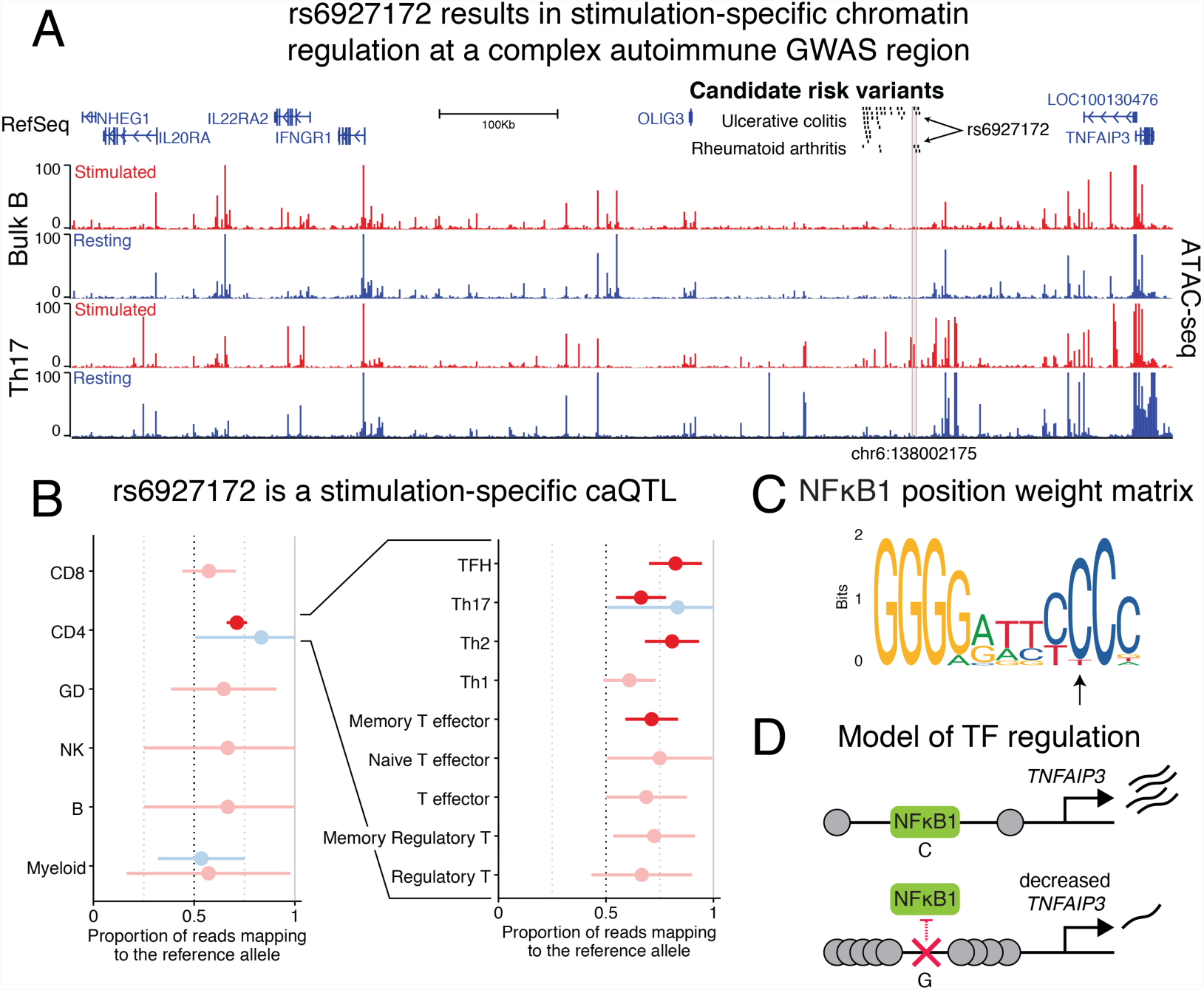
Identifying rs6927172 as a stimulation-specific chromatin regulator in a complex autoimmune GWAS region. (A) Chromatin accessibility profile for stimulated (red) and resting (blue), bulk B (top) and Th17 (bottom) cells, around variant rs6927172. This region contains significant GWAS signals for ulcerative colitis and rheumatoid arthritis, but the causal variant(s) have not been determined (credible set indicated). (B) Allele-specific ATAC-seq reads at rs6927172 in the three heterozygous donors (the fourth was not heterozygous at this site). Displayed is the proportion of reads mapping to the reference allele. Error bars represent 98% confidence intervals and were computed from read depth. Significant (p < 0.01) allelic imbalance associations are colored.(C) The canonical PWM for the p50 subunit of NFkB1, as downloaded from the Jaspar TF motif database. The heterozygous allele disrupts the nucleotide indicated by the arrow. (D) A proposed negative feedback model of gene regulation linking NFkB1 to *TFNAIP3*.

## Discussion

While increasing evidence points to an important role of stimulation response in autoimmune dysregulation, little work has been done to characterize shared and cell subset-specific stimulation effects across multiple differentiation lineages. Therefore, we developed a map of stimulation-responsive chromatin accessibility and transcription in a diverse set of purified immune cell populations. Using these data, we characterized the contributions of cell type and stimulation states to the regulation of chromatin accessibility. We found that stimulation-associated changes cause dramatic chromatin remodeling including a shared stimulation response in B and T cells, driven in part by activation of BATF.

To study the impact of genetic variation on the chromatin landscape, we used allele-specific measurement of open chromatin to identify genotype-dependent sites (ASC). While the majority of ASC sites are shared between closely related cell types, only 60% of ASC sites are shared for comparisons between lineages and between stimulated and resting cells. Most non-sharing of ASC sites is due to differential accessibility between cell types/conditions, while a smaller fraction is due to differential regulation of shared open sites as exemplified by BATF activity in stimulated cells.

Several past studies have mapped GWAS signals to cell type-specific functional elements, allowing the prediction of the cell types that may be involved in disease pathology (*3-8, 40*). We observed significant autoimmune trait heritability within accessible regions of distinct stimulated cell types, suggesting contributions of multiple stimulation-response components to autoimmunity. Moreover, we showed that genetic variation associated with stimulation-specific gene regulation is significantly underrepresented in existing large-scale eQTL datasets from whole blood PBMCs.

As a concrete example of this point we identified rs6927172, which was associated with CD4^+^ T cell stimulation-specific changes in chromatin accessibility near *TNFAIP3* and affects the predicted binding affinity of the p50 subunit of NF?B1. This region has been associated with several autoimmune phenotypes, such as variable patient responses to anti-TNF treatment (*41*), ulcerative colitis (*17*), and rheumatoid arthritis (*42*). Interestingly, the variant was not detected as an eQTL in GTEx v7, likely because tissues represent cell mixtures with generally low proportions of immune cells and even lower proportions in activated states. However, Wu et al. demonstrated that the disruption of an 11bp region harboring rs6927172 significantly decreased gene expression of *TNFAIP3* in stimulated HEK293 T cells (*43*), suggesting this variant drives *TNFAIP3* expression. Under resting state, T cells constitutively express *TNFAIP3* to suppress the activity of stimulation-responsive signaling, in part by deactivating NFkB1 activity. Upon stimulation, Coornaert et al. found that *TNFAIP3* expression first decreases and then reappears, suggesting that the initial removal of A20 (encoded by *TNFAIP3)* is essential for optimal NFkB1 activation (*44*). While the mechanism that leads to the opening of the region containing rs6927172 and the suppression of A20 is unclear, we speculate a model whereby A20 is first down-regulated by other factors, allowing activation of NFkB1. Next, NFkB1 binds to the region containing rs6927162, resulting in the reappearance of A20 expression. Thus, rs6927172 likely prevents the return of A20 by disrupting the binding of NFkB1, which subsequently results in inappropriate NFkB1 signaling.

Taken together, these data provide insights into the interplay of specific loci and regulatory networks involved in the human immune system. Additionally, we present a powerful platform to map genetic variants to cell type and context-specific functional regions genome-wide, and therefore, to their context-specific effects on disease phenotypes.

## Methods

### Data collection

#### Sample collection and processing

This study was approved by the UCSF Committee on Human Research and written consent was obtained from all donors. Peripheral blood mononuclear cells (PBMCs) were isolated from whole blood (~450 ml) using Ficoll-Plaque Plus (GE Healthcare, IL, USA) centrifugation. Bulk population of CD4^+^ T cells, B cells and monocytes were positively enriched, whereas pan T cells, NK cells and dendritic cells were negatively enriched using magnetic beads prior to sorting (STEMCELL, Canada). Different cell subsets were sorted on a FACS Aria flow cytometer (BD Biosciences, CA, USA) up to >95% purity (Table S1). Sorted cells were washed once in PBS, cryopreserved in Bambanker freezing media (LYMPHOTEC Inc, Japan) for ATAC experiments and in TriReagent (Sigma-Aldrich, MO, USA) for RNA experiments. Cells frozen in Bambanker freezing media were stored in liquid nitrogen until ready to use. Cells frozen in TriReagent were stored at −80 °Cuntil further use.

#### Fetal sample processing

Human thymus was obtained from 18-to 22-gestational-week specimens under the guidelines of the Committee on Human Research (UCSF IRB)–approved protocols from the Department of Obstetrics, Gynecology and Reproductive Science, San Francisco General Hospital. Fetal samples were obtained after legal, elective termination of pregnancy with written informed consent for fetal tissue donation to biomedical research. Consent for tissue donation was obtained by clinical staff after the decision to pursue termination was reached by patients. Personal Health Information and Medical Record Identifiers/access is at no point available to researchers, and no such information is associated with tissue samples at any point. Tissue was washed and cut into small pieces using scissors. Thymocytes were extracted by mashing tissue pieces gently using the back of a sterile syringe. To extract TECs, remaining tissue pieces were digested for 30 min at 37 °using medium containing 100 μg ml^-1^ DNase I (Roche, Switzerland) and 100 μg ml^-1^ Liberase TM (Sigma-Aldrich, MO, USA) in RPMI. Fragments were triturated through a 5-ml pipette every 6 min to mechanically aid digestion. At 30 min, tubes were spun briefly to pellet undigested fragments and the supernatant was discarded. Fresh digestion medium was added to remaining fragments and the digestion was repeated using a glass Pasteur pipette for trituration. Supernatant from this second round of digestion was also discarded. A third round of enzymatic digestion was performed using digestion medium supplemented with trypsin-EDTA for a final concentration of 0.05%. Remaining thymic fragments were digested for another 30 min or until a single cell suspension was obtained. The cells were moved to cold MACS buffer (0.5% BSA, 2 mM EDTA in PBS) to stop the enzymatic digestion. Following digestion, TECs were enriched by density centrifugation over a three-layer Percoll gradient with specific gravities of 1.115, 1.065 and 1.0. Stromal cells isolated from the Percoll-light fraction (between the 1.065 and 1.0 layers) were washed in MACS buffer. Samples were sorted on FACS Aria flow cytometer (BD Biosciences, CA, USA) up to >95% purity. Sorted cells were washed once in PBS, cryopreserved in Bambanker freezing media (LYMPHOTEC Inc, Japan) for ATAC experiments and in TriReagent (Sigma-Aldrich, MO, USA) for RNA experiments. Cells frozen in Bambanker freezing media were stored in liquid nitrogen until ready to use. Cells frozen in TriReagent were stored at −80°Cntil further use.

#### Ex vivo activation

Freshly sorted cells were cultured in RPMI-1640 medium supplemented with glutamine, sodium pyruvate, penicillin, streptomycin, non-essential amino acid and 10% FCS (Sigma-Aldrich, MO, USA). T lymphocytes were stimulated for 24 hours with anti-human CD3/CD28 dynabeads (Thermo Fisher Scientific, MA, USA) at a 1:1 cell to bead ratio and human IL-2 (UCSF Pharmacy, 300 unit/ml for regulatory T cells, 50 unit/ml for other T lymphocytes). B lymphocytes were activated for 24 hours with 10 *μ*g/ml F(ab)‘2 anti-human IgG/IgM (Affymetrix, CA, USA) and 20 ng/ml human IL-4 (Cell Sciences, MA, USA). NK cells were activated using 2 conditions: 1) with the NK Cell Activation/Expansion Kit as per manufacturer’s instructions for 48 hours (Milteny, Germany) and 2) with 500 unit human IL-2 for 24 hours. Monocytes were stimulated with 100 ng/ml and 1*μ*g/ml LPS (Sigma-Aldrich, MO, USA) for 6 hours and 24 hours, respectively. The detailed activation conditions are listed in Table S1). After stimulation, cells were washed once with PBS, cryopreserved in Bambanker freezing media and TriReagents, and stored in liquid nitrogen until ready to use.

#### RNA-seq library preparation

Cells frozen in TriReagent were thawed at room temperature for 10 min. Technical replicates were done for each cell aliquot. One hundred *μ*l chloroform was added to 500 *μ*l of sample, mixed, incubated at room temperature for 10 min and centrifuged at 12,000 g for 10 min at 4°C. The aqueous phase was transferred to a new tube and an equal volume of 100% ethanol was added. For further RNA extraction the Direct-zol RNA MicroPrep kit (Zymo Research, CA, USA) was used. Samples were mixed thoroughly before transferring onto a Zymo-Spin IC Column and RNA was extracted according to the manufacturer’s protocol starting at step 2. RNA sequencing library preparation was based on the Smart-seq2 protocol (*45*). For oligo dT annealing, 2 *μ*l RNA was mixed with 1 *μ*l recombinant RNase inhibitor (Takara Bio USA, CA, USA), 1 *μ*l of 10 *μ*M oligo dT (Integrated DNA Technologies, [IDT], IA, USA) and 1 *μ*l of 10 mM dNTPs (New England Biolabs, MA, USA), and incubated at 72°C for 3 min. Reverse transcription and template switching were done in 1x first strand synthesis buffer (Thermo Fisher Scientific, MA, USA), 100 U SuperScript II reverse transcriptase (Thermo Fisher Scientific), 10 U RNase inhibitor (Takara Bio USA), 5 mM DTT, 1 M Betaine (Sigma-Aldrich, MO, USA), 6 mM MgCl_2_, and 1 *μ*M TSO (IDT). The mixture was incubated at 42°C for 90 min, 10 cycles of 50°C for 2 min and 42°C for 2 min, followed by a final incubation of 15 min at 70°C. Transcribed RNA was amplified in 1x KAPA HiFi HotStart ReadyMix (Kapa Biosystems, MA, USA), 0.1 *μ*M IS primer and 0.6 x Sybr Green (Thermo Fisher Scientific) as follows: 98°C for 3 min, 18 cycles of 98°C for 20 seconds, 67°C for 15 seconds, followed by a final extension at 72°C for 5 min. Samples were purified using AMPure XP beads (Beckman Coulter, CA, USA) in a 1:1 ratio. Tagmentation of amplified cDNA was carried out using the Nextera XT DNA library kit (Illumina, CA, USA). In a 20 *μ*l reaction, 150-200 pg cDNA was added to 1x TD buffer (Illumina) and 5 *μ*l of ATM (Illumina). The mixture was incubated at 55°C for 5 min. Five *μ*l of NT buffer was added to the mixture to stop the fragmentation reaction and incubated for 5 min at room temperature. Tagmented libraries were further amplified in a total reaction volume of 50 *μ*l by adding 15 *μ*l of NPM (Illumina) and 1.25 *μ*M of each, i7 and i5, custom Nextera barcoded PCR primers (*46*) for 3 min at 72°C, 30 seconds at 95°C, and 12 cycles of 10 seconds at 95°C, 30 seconds at 55°C, and 30 seconds at 72°C. Amplified libraries were purified using AMPure XP beads (Beckman Coulter) using a 1:1 ratio.

#### ATAC-seq library preparation

ATAC-seq libraries were prepared following the Fast-ATAC protocol (*20*). In detail, frozen cells were thawed in a water bath, resuspended in 2 ml media and centrifuged at 500 g for 5 min. Pelleted cells were resuspended in 1 ml and counted using a hemocytometer. For the ATAC reaction, we took aliquots of 10,000 live cells, unless limited by cell number. Technical replicates were done for all samples. Cells were spun down at 4°C at 500 g for 5 min, washed once in cold PBS and resuspended in the Tn5 reaction buffer (1x TD buffer [Illumina, CA, USA], 50 *μ*l TDE1 Tn5 transposase [Illumina] per ml Tn5 reaction buffer and 0.01% Digitonin). 5,000-10,000 cells were transposed in 50 *μ*l of reaction buffer. The reaction volume of samples containing fewer than 5,000 cells was linearly scaled to the number of cells present whereby, for example, 4,000 cells were done in a 40 *μ*l reaction and 2,500 cells were done in a 25 *μ*l reaction. The Tn5 reaction mix was incubated at 37°C for 30 min at 300 rpm. Transposed samples were purified using MinElute PCR purification columns according to the manufacturer’s protocol (Qiagen, Germany). Purified samples were amplified and indexed using custom Nextera barcoded PCR primers (Table S6) as described in 21. Amplified libraries were purified using MinElute PCR purification columns.

#### DNA sequencing

To circumvent index hopping, subpools of samples were created, ran over a 2.5% Agarose gel and size selected, excluding high molecular weight DNA, free PCR primers, and primer dimers. Samples were purified using a MinElute Gel Purification kit according to manufacturer’s instructions (Qiagen). Prior to sequencing, samples were amplified in 1x NEBNext Master Mix (New England BioLabs, MA, USA), 1.25 *μ*M oligo C (Illumina P5) and 1.25 *μ*M oligo D (Illumina P7) using the following cycle conditions: 30 seconds at 98°C followed by 3-4 cycles of 10 seconds at 98°C, 30 seconds at 63°C and 1 min at 72°C. All samples were purified using MinElute PCR purification columns (Qiagen). Samples run on the NovaSeq 6000 were additionally purified using 1x AMPure XP beads (Beckman Coulter, CA, USA) to remove free oligo C and D primers. ATAC-seq samples were sequenced on a HiSeq 4000 in paired-end 76 bp cycle mode or on a NextSeq 500, without prior gel clean-up, in paired-end 76 bp cycle, high-output mode. RNA-seq (cDNA) libraries were either sequenced on a NovaSeq using a S2 flow cell in paired-end 100 bp cycle mode or on a HiSeq 4000 in paired-end 76 bp cycle mode.

#### Genotyping

Three hundred *μ*l of donor blood (donors: 1001,1002, 1003 and 1004) was used to extract genomic DNA using DNeasy Blood and Tissue Kits (Qiagen, Germany) as per manufacturer’s protocol. We genotyped 958,497 markers using Infinium OmniExpressExome-8 v1.4 kit (Illumina, CA, USA). We phased and imputed all biallelic common variants (MAF > 1%) using the Michigan imputation server that ran minimac3 with 1000G Phase 3 v5 as the reference genome and Eagle v2.3 (*47*). All genotypes with an imputed heterozygous probability of 0.9 or greater were included in downstream allele-specific chromatin accessibility analysis.

### Collection of publicly available data

#### Progenitor RNA-seq and ATAC-seq data from GEO

ATAC-seq and RNA-seq of hematopoiesis progenitors and several differentiated cell types were downloaded (*20*), and processed through the respective ATAC-seq and RNA-seq pipeline described below. Only data from healthy controls was included throughout this study.

#### ATAC-seq data from ENCODE

To serve as a negative control for GWAS enrichment analyses, we collected data from ATAC-seq samples from tissues with low proportions of immune cells. We used the ENCODE data portal to download all available raw fastq ATAC-seq files from the calf muscle and breast epithelium human tissues. All data were processed with the same ATAC-seq data processing pipeline described below.

#### Obtaining GWAS summary statistics

We downloaded the full set of GWAS summary statistics of 13 complex traits from 9 autoimmune traits and 4 primarily non-autoimmune and thus negative control traits (Sunburn, Alzheimer’s disease, Type 2 diabetes, and Schizophrenia) that have been previously aggregated (*48*). For analyses that relied on fine-mapped disease-associated variants, we downloaded the list of PIC variants. These represent individual GWAS regions that have been fine-mapped with a previously described statistical method (*4*).

#### Obtaining Blood eQTL summary statistics

From the GTEx data portal we downloaded v7 eQTL estimates for all SNP-gene pairs tested with whole blood gene expression.

### Data analysis

#### RNA-seq

We trimmed remaining transposase adapters with cutadapt version 1.13 with a –minimum-length of 20 and an –overlap of 5 in paired-end mode (*49*). RNA-seq reads were pseudoaligned using Kallisto version 0.43.0 (*50*). The Kallisto index was made with default parameters and the gencode v25lift37 fasta file (*51*) and was run in quant mode with default parameters. Following pseudoalignment we computed gene abundances using tximport version 1.2.0 (*52*). We excluded donor cell type samples with fewer than one million total reads from all technical replicates or that were extreme PCA outliers. Discarded cell types include thymocyte subsets and TEC cells.

#### ATAC-seq

We trimmed remaining transposase adapters with cutadapt version 1.13 with a –minimum-length of 20 and an –overlap of 5 in paired-end mode (*49*). We aligned ATAC-seq reads using bowtie2 version 2.2.9 with default parameters and a maximum paired-end insert distance of 2kbps. The bowtie2 index was constructed with the default parameters for the hg19 reference genome. We filtered out reads that mapped to chrM and used samtools version 1.4 to filter out reads with MAPQ < 30 and with the flags ‘-F 1804’ and ‘-f 2’. Additionally, duplicate reads were discarded using picard version 1.134 (*53*). We calculated percentage aligning to mitochondria, percentage within blacklist regions (*54*), and enrichment of reads at TSS regions relative to 2kb away using the RefSeq gene annotation (referred to elsewhere as the TSS enrichment). Chromatin accessibility peaks were identified with MACS2 version 2.1.1 under default parameters and ‘--nomodel --nolambda --keep-dup all --call-summits’. The count of absolute peaks per cell type refers to the number of peak regions reported in the ‘narrowPeak’ file (peaks with multiple summits are only counted once). We reported the peak count estimate after linearly adjusting for estimated confounding effects from sample read depth, and TSS enrichment (a proxy for sample quality). A consensus set of peaks was defined by merging overlapping (1bp or more) peaks identified in at least two samples across all samples. This set of peaks had a median peak length of 346 bps following removal of peaks that were greater than 3 kb or non-autosomal. We then used the ‘get_count’ function from the nucleoATAC python package to count the number of fragments within the consensus peak set across all samples (*55*). We excluded donor samples with fewer than 5 million reads from all technical replicates that passed quality filters, TSS enrichment score less than 4, more than 0.5% of reads mapping to a known set of region blacklist (*54*), more than 25% of reads mapping to the mitochondria, or that were extreme PCA outliers. We used deeptools version 2.5.1 to generate bigwig tracks with a genomic bin size of 10 bp, RPKM normalization and the ‘—extendReads’ parameter.

#### Exploratory analysis

We used tSNE, PCA, and k-means clustering to explore trends in gene expression and chromatin accessibility variation. As input to these methods, we used the read count matrices corrected for sample quality (with TSS enrichment as a proxy) and batch effects (with donor as a proxy). After using trimmed mean of M-values (TMM) to estimate scaling factors (*56*) and applying the voom transformation (*57*), we used the ‘removeBatchEffect’ function from limma to regress out batch and sample quality effects (*58*). Aside from the removal of these effects, normalized counts are equivalent to the addition of a 0.5 pseudocount to the fragment count and a log2 transformation of the fragment counts per million (CPM). For analyses that included previously published samples, we did not remove batch effects, because these batches do not have overlapping cell types. However, batch appeared to have a minimal effect on sample clusters and these analyses were exploratory in nature. The package Rtsne version 0.13 was used for tSNE analysis with default parameters, unless there were too few samples in which case the perplexity was set to 10 (*59*). Additionally, we performed k-means clustering (with k set to the total number of cell-type by condition pairs) and then estimated the accuracy of ATAC-seq and RNA-seq unsupervised clustering by computing the HA-adjusted RAND index (*60*) considering the known cell-type/condition pairs as the ground truth clustering. We repeated the clustering 100 times to estimate the average HA-adjusted RAND index. When comparing HA-adjusted RAND index values between the RNA-seq and ATAC-seq samples, we used the intersection of samples.

#### Differentially expressed genes

For this analysis, we included only unique protein coding genes from gencode v25. Additionally, we eliminated all genes that had below ten counts per million (CPM) in at least two biological replicate samples, resulting in 13,512 genes tested. We used the TMM normalization method to compute scale factors for each sample, and voom to compute weights capturing the relationship between gene expression mean and variance. For each cell type, we used limma to estimate the log2 fold change (log2FC) of gene expression for each gene upon stimulation. We included donor in the design matrix to correct for donor-specific effects. We considered a gene differentially expressed if the limma-reported Benjamini-Hochberg (BH) corrected *p*-value (*61*), which we refer to as a *q*-value, was less than 0.01 and the absolute log2FC was greater than 1.

#### Differentially accessible chromatin regions

Starting with the consensus peak set we eliminated all peaks with fewer than one CPM in at least two biological replicate samples, resulting in 671,448 tested peaks. We used the TMM method to compute scale factors for each sample, and voom to compute weights capturing the relationship between mean and variance chromatin accessibility across samples. For each cell type we used limma to estimate the log2FC of each chromatin accessibility region upon stimulation, while controlling for donor and the enrichment of reads at TSSs (the latter serves as a proxy for sample quality). We considered a region a significant differentially accessible chromatin peak if the limma-reported BH corrected *p*-value was less than 0.01 and the estimated absolute log2FC from the resting to stimulation condition was greater than 1. We initially considered different stimulation conditions of NK and monocyte samples separately. However, estimated log2FC stimulation effects were nearly identical among significant stimulation peaks when the conditions were pooled or unpooled (adjusted R^2^ = 0.88 for Mature NK cells and R^2^ = 0.99 for Monocytes). Thus, we reported summary statistics from the pooled differential accessibility analysis.

#### Quantifying the amount of shared stimulation response

We aimed to quantify sharing of simulation effects between two cell types, denoted A and B. To do this we first determined the set of differentially accessible peaks (q<0.01 and |log2FC|>1.0) from cell type A (using limma-voom, see *Differentially accessible chromatin regions* above). For this set of peaks, we computed Pearson correlation (with the ‘cor’ function in R) between stimulation effects (log2FC) in cell type A and B. This analysis results in asymmetric sharing estimates conditioning on significant peaks from cell type A or B. We found that the asymmetric estimates were broadly consistent, so we report the mean correlation weighted by the number of significant peaks in each cell type (individual asymmetric correlation estimates can be found in Table S7).1

#### Allele-specific chromatin accessibility

We collected all aligned reads that overlap heterozygous sites. Initially, we used the set of heterozygous sites identified with ‘HaplotypeCaller’ from GATK version 3.7 with ‘—minPruning 10’ and ‘-stand_call_conf 20’ run on a donor-specific bam file including all samples from the donor (*62*). The final set of heterozygous sites that we used for analysis were the intersection of the GATK and genotyped sites passing filters (see *Genotyping* above). Next, we passed all aligned reads that overlap heterozygous sites through WASP filtering (*34*). Briefly, the read is remapped with the SNP allele flipped (we only examine biallelic sites) and is only retained if it maps to the same location. This filtering approach effectively eliminates reference genome biases enabling unbiased estimation of the proportion of reads mapping to the reference allele. At each heterozygous site we counted the number of WASP filtered reads mapping to the reference and the alternative allele. We computed a p-value per heterozygous site using a binomial test, and for each sample corrected for multiple hypotheses by computing BH q-values. P-values were converted to BH q-values using the ‘p.adjust’ function in R.

#### Estimating the proportion of shared allele-specific imbalance effects

We first computed the posterior mean and variance of the proportion of reads mapping to the reference allele, assuming a uniform prior on the proportion and a binomial likelihood. The posterior under this model is Beta(*r* + 1, *a* + 1), where r and a are the numbers of reads mapping to the reference and alternative allele respectively. At each heterozygous site, we defined the effect as the difference between the proportion of reads mapping to the reference and the expectation under the null of no allele-specific effect, i.e., 50%.

We aimed to estimate the proportion of shared imbalance effects between two cell types, denoted A and B. To do this we first determined the set of significant ASC sites (q < 0.01) from cell type A (see *Allele-specific chromatin accessibility* above). We collected effects and variances for these ASC sites in cell type B, and then used ashR (*63*) to estimate the proportion of these effects that are nonzero. While comparable methods such as Storey’s pi1 only uses *p*-values (*64*), ashR leverages both effect size and standard error estimates to improve power. We reported the average sharing estimates, however individual values can be found in Table S8.

#### Variance decomposition

We were interested in decomposing the total variance of chromatin accessibility across samples into variance components attributable to specific factors. Therefore, we fitted a random effects model:

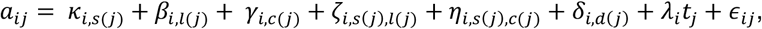

where chromatin accessibility (*a*) at peak *i*for a sample *j* is a function of the effects of the stimulation condition (*κ*), lineage (*β*), cell type (*γ*), lineage/stimulation interaction (*ζ*), cell/stimulation interaction (*η*), donor (*δ*), TSS enrichment (*t*) of ATAC-seq reads (*λ*), and the residual error (*ε*). For notational convenience, we define a function for each feature in the model that looks up sample-specific information, i.e., *s*(*j*) represents the stimulation condition associated with sample *j*. We represented accessibility with the log2(CPM) ATAC-seq read counts (with the addition of a pseudocount of 0.5) at consensus peaks across samples, which were normalized for read depth with TMM normalization. Additionally, we scaled accessibility at each peak across samples to have mean=0 and variance=1. We included the effects of *d* and *t* (as a proxy for sample quality), to control for their effects, since the other parameters are our primary interest. Across all peaks, we modelled the distribution of effects:

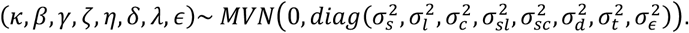

We used a maximum likelihood approach to jointly estimate the *σ*2 parameters for each factor. To obtain robust estimates we found it beneficial to pool peaks. We found pooling 100 peaks represented a good compromise between computational cost and statistical robustness. To assess uncertainty of variance estimates, we repeated the analysis on 100 sets of 100 randomly selected peaks with replacement. The total biological variance explained (TBVE) by the factors of interest is,

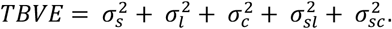

Therefore, the proportion of biological variance explained (PBVE) contributed by a factor is the variance estimate *σ*2 for that factor divided by t. We listed the median value across all bootstrap replicates. For results reported we limited our analysis to cell types from the four donors with the most cell samples collected and excluded cell types with fewer than three biological replicates.

#### TF position weight matrix (PWM) motif analyses

For determining PWMs enriched in open chromatin regions we used chromVAR version 1.0.1 with default parameters on read counts within consensus peaks of samples merged by donor (*65*). Following identification of condition-associated TFs with chromVAR we wanted to examine the effect of a few of these TFs on allele-specific chromatin accessibility. We used the PWM of a TF of interest to predict the binding affinity of a 41 bp genomic region centered on the heterozygous site. The binding affinity or match score was computed using the ‘motifmatchr’ R package (*66*), which is a wrapper for the MOODS motif matching suite (*67*). The relative binding score was determined by subtracting the binding affinity match score of the alternative allele from that of the reference allele. As a threshold for presence or absence of motif matching we used a p value cutoff of 3 × 10^-3^. In this way we grouped heterozygous sites into three groups: predicted TF affinity for the reference (relative match > 1), alternative (relative match < −1) or no preference (absolute value of the relative match < 0.01).

#### GWAS and eQTL enrichments

To identify genomic annotations enriched for genetic trait heritability we used LD Score regression version 1.0.0 under the partitioning heritability mode with default parameters. We excluded SNPs from the MHC region for this analysis. To assess enrichment of RA signal in ASC sets, we compared against the distribution of the p-values for the RA GWAS genome-wide (the same summary statistics that were used in the LD Score analysis). To asses enrichment of blood eQTL signal in ASC sets, we compared against the distribution of p-values from all variant-gene pairs available.

#### Peak clustering

To test whether disease heritability was enriched within stimulation-specific chromatin accessible from B and T cell lineages we used a supervised peak clustering approach. First, we scaled the matrix of ATAC-seq read counts per sample (indicated by *j*) across all consensus peaks (indicated by t) to values between 0 and 1 with

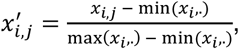

where *x*’ represents the scaled matrix. Peaks in a sample that were in the top decile were automatically set to 1 to represent the fully accessible state. Per peak we computed the median scaled accessibility across samples from the same broad cell type and condition.

Our goal was to identify peaks that express a specific accessibility profile. We defined a profile of interest with a vector of length equal to the number of merged lineage and condition samples with values of either 0 or 1 corresponding to closed or open chromatin accessibility. We consider 11 profiles of interest (Figure S5D). To identify peaks with a similar profile we computed the average Euclidean distance between each of the ideal accessibility profiles and each peak. Peaks more similar to an ideal peak profile should have smaller distances to the peak profile. Additionally, when computing the distance, we incorporated a weight per sample to influence the importance of matching accessibility in different merged samples. This was important to find peaks that were accessible in resting samples (weight of 1), while allowing for the possibility that the peak was accessible in the same lineage but stimulated samples (weight of 0).

To determine a distance cutoff of a peak cluster, we permuted the peak accessibility values for each sample and computed a null distribution of distances for each lineage and peak cluster type. We used a peak distance threshold resulting in fewer than 5% false positives. Finally, peaks passing this peak distance threshold were assigned to a single profile of interest based on the minimum distance, thus forming disjoint sets of accessible regions.

## Data availability

Our GEO RNA-seq data repository: GSE118165

Our GEO ATAC-seq data repository: GSE118189

Progenitor data GEO accession: GSE74912

## URLs

ENCODE data portal: https://www.encodeproject.org/

GTEx data portal: https://www.gtexportal.org/home/datasets

GWAS summary statistics: https://data.broadinstitute.org/alkesgroup/SLDP/sumstats/complex/

## Acknowledgements

We thank D. Yao for helping process samples, Chun Jimmie Ye and members of the Greenleaf, Marson, and Pritchard laboratories for helpful conversations and manuscript feedback. We relied on the Flow Cytometry Core at UCSF, which was supported by the Diabetes Research Center grants NIH P30 DK063720 and 1S10OD021822-01, and sequencing data generated on an Illumina HiSeq 4000 that was purchased with funds from NIH under award number S10OD018220. Sequencing that was generated on an Illumina NovaSeq was supported by the Chan Zuckerberg Biohub. Some of the computing for this project was performed on the Sherlock cluster. We would like to thank Stanford University and the Stanford Research Computing Center for providing computational resources and support that contributed to these research results. Support for D.C. was provided by NLM Training Grant Number T15LM007033. A. Mezger is supported by the Swedish Research Council (grant 2015-06403). This work was supported by NIH grants (1R01HG008140, P30AR070155 to L.A.C., DP3DK111914-01 to A. Marson, P50HG007735 to W.J.G., UM1HG009442 to W.J.G, U19AI057266 to W.J.G.), the Howard Hughes Medical Institute (J.K.P.), the Rheumatology Research Foundation (L.A.C.), the UCSF-Stanford Arthritis Center of Excellence (L.A.C.)(supported in part by the Arthritis Foundation), the Rita Allen Foundation (W.J.G.), the Human Frontiers Science Program grant RGY006S (W.J.G.), the Burroughs Wellcome Fund (A. Marson), National Multiple Sclerosis Society (A. Marson; CA 1074-A-21), and the Chan Zuckerberg Biohub (W.J.G., A. Marson).

## Author contribution

Conceptualization, D.C., M.L.T.N., A. Mezger, L.A.C., W.J.G., A. Marson, and J.K.P.; Investigation, M.L.T.N., A. Mezger, A.K., V.N., N.L., B.W., J.T., and A.V.P.; Formal Analysis, D.C., D.A.K., Z.G., andJ.V.R.; Resources, M.S.A., T.D.B, W.J.G., A. Marson, and J.K.P.; Funding Acquisition, L.A.C., W.J.G.,A. Marson, and J.K.P.; Writing – Original Draft, D.C., M.L.T.N., and A. Mezger; Writing – Review & Editing, D.C., M.L.T.N., A. Mezger, L.A.C., W.J.G., A. Marson, and J.K.P.; Supervision, W.J.G., A. Marson, and J.K.P.

---

**Figure S1.**
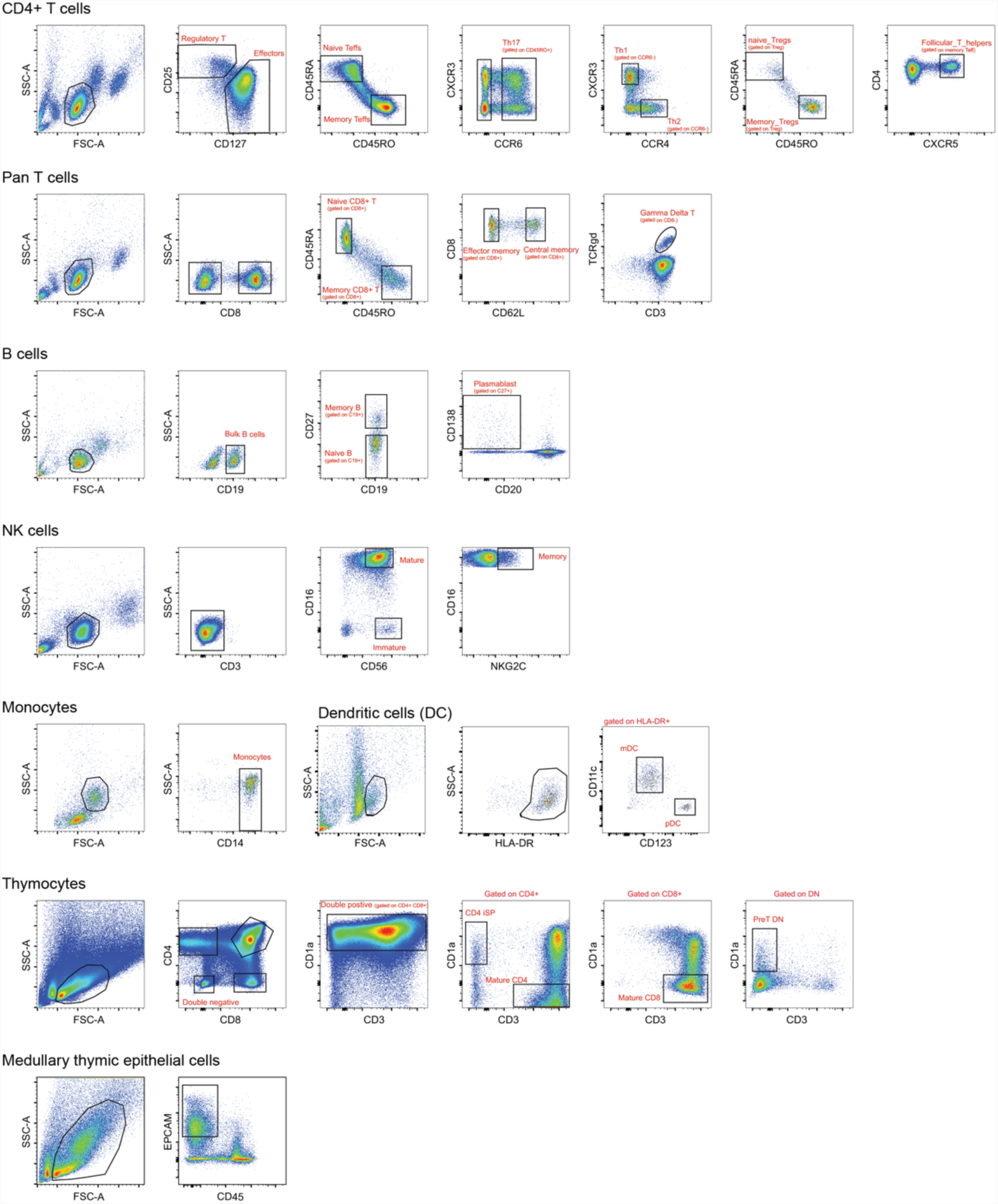
Gating strategies and surface markers for cell sorting. Representative flow cytometry plots of the sorting panels for each population of immune cells. Monocytes, bulk CD4+ T and B cells were positive enriched with magnetic beads prior to sorting. Pan T cells, DCs and NK cells were negatively enriched.

**Figure S2.**
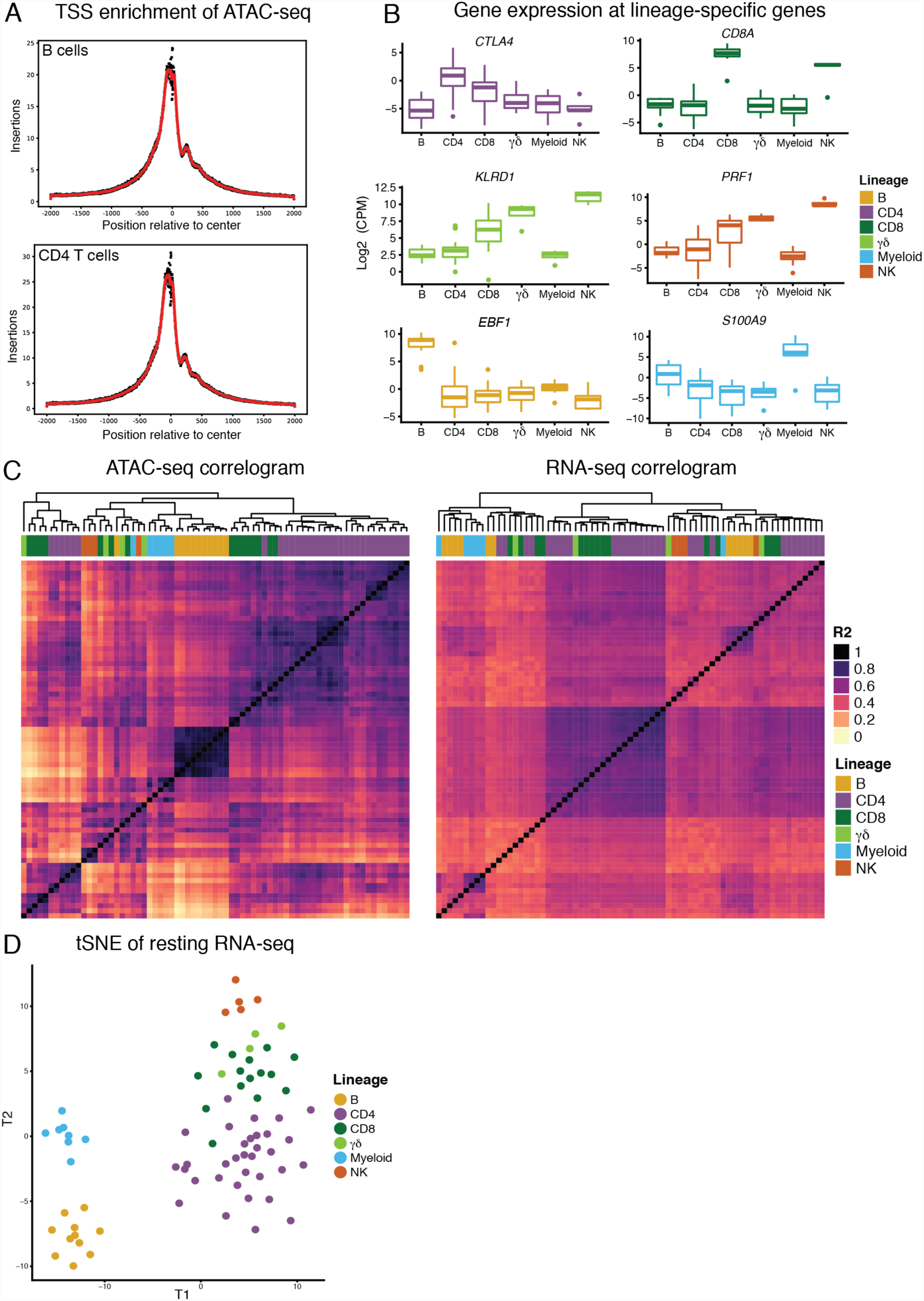
Overview of ATAC-seq and RNA-seq data. (A) Enrichment of ATAC-seq reads relative to RefSeq TSSs for B cells (top) and CD4^+^ T cells (bottom). (B) Examples of gene expression across lineages, y-axis represents log2 (count per million (CPM)). (C) Correlogram of Pearson’s R2 for ATAC (left) and RNA (right) for all overlapping samples. (D) RNA-seq tSNE of batch corrected read count matrix at protein coding genes.

**Figure S3.**
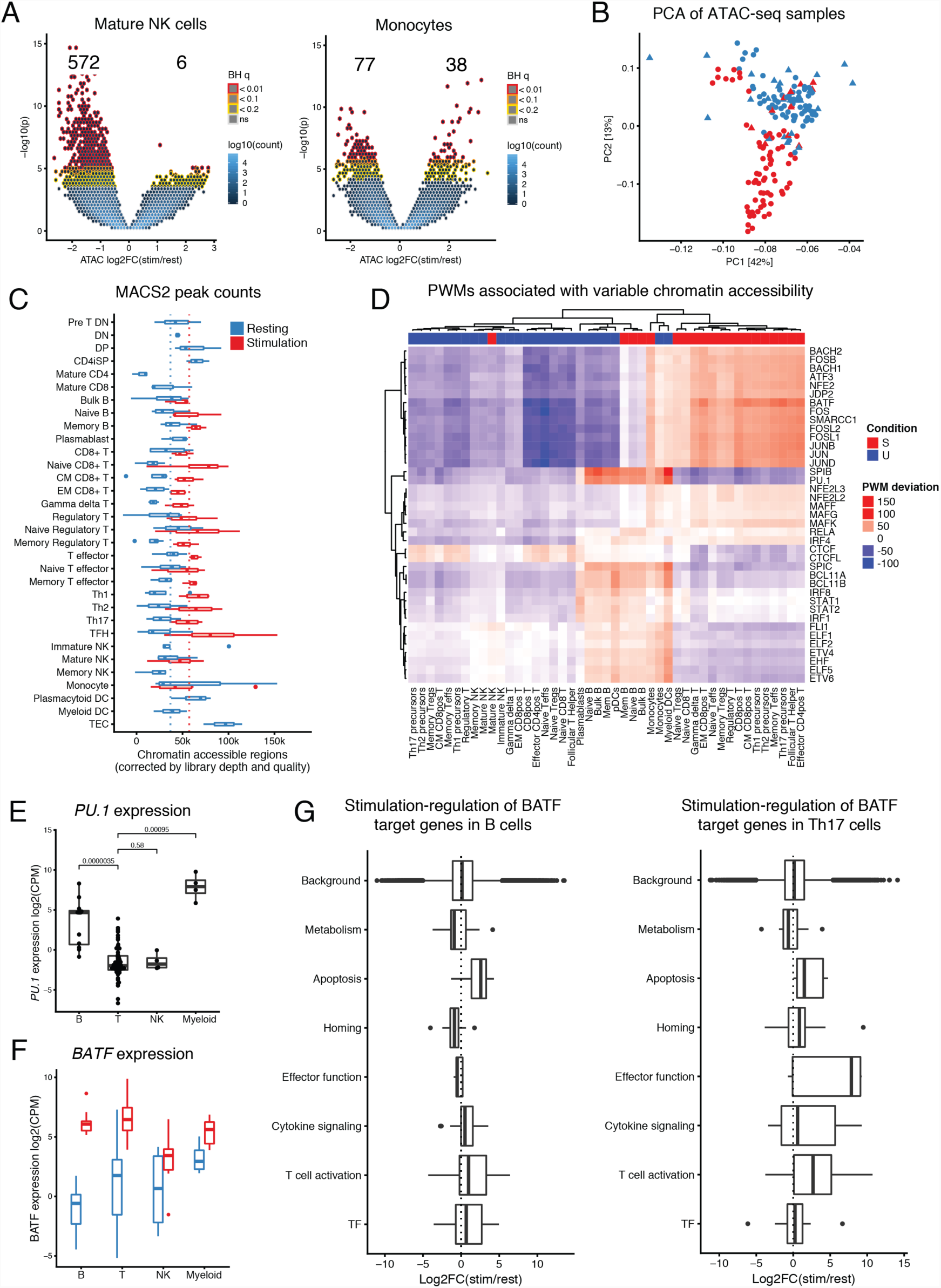
Characterization of ATAC peaks and gene expression. (A) Volcano plot of differentially accessible chromatin regions for mature NK (left) and monocytes (right). (B) PCA of resting (blue) and stimulated (red) ATAC-seq samples including the innate cells (triangles). (C) MACS2 peak count across different cell types stratified by condition. MACS2 peak counts were normalized for read count and sample quality effects. The dotted lines indicate the median count of accessible regions in stimulated (red) and resting (blue) samples. (D) Heatmap of chromVar TF binding deviation scores for several PWMs (y-axis) and samples (x-axis). (E) Gene expression of *PU.1* separated by lineage.(F) Expression of *BATF* grouped by lineage and condition. Resting samples indicated in blue and stimulated samples in red. (G) Expression of BATF target genes in B cells (left) and Th17 precursors (right).

**Figure S4.**
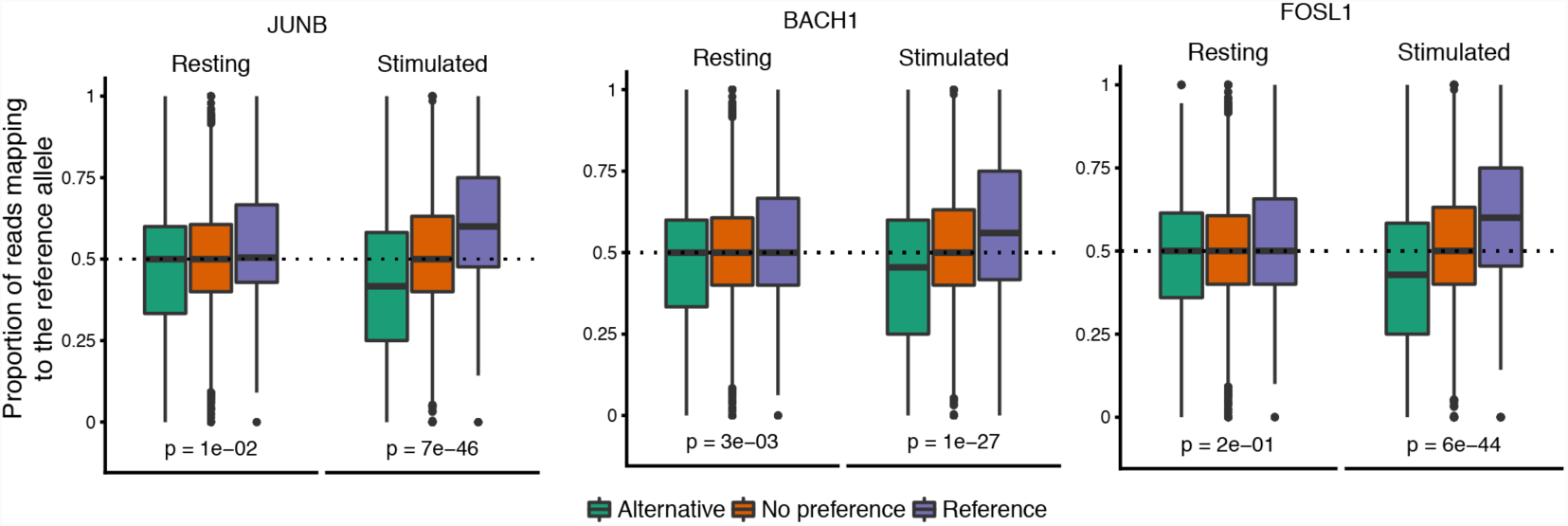
Stimulation-specific chromatin imbalance in Th1 precursor cells. Changes in the proportion of reads mapping to the reference allele at heterozygous sites as a function of stimulation condition and PWM-predicted JUNB, BACH1 and FOSL1 binding affinity.

**Figure S5.**
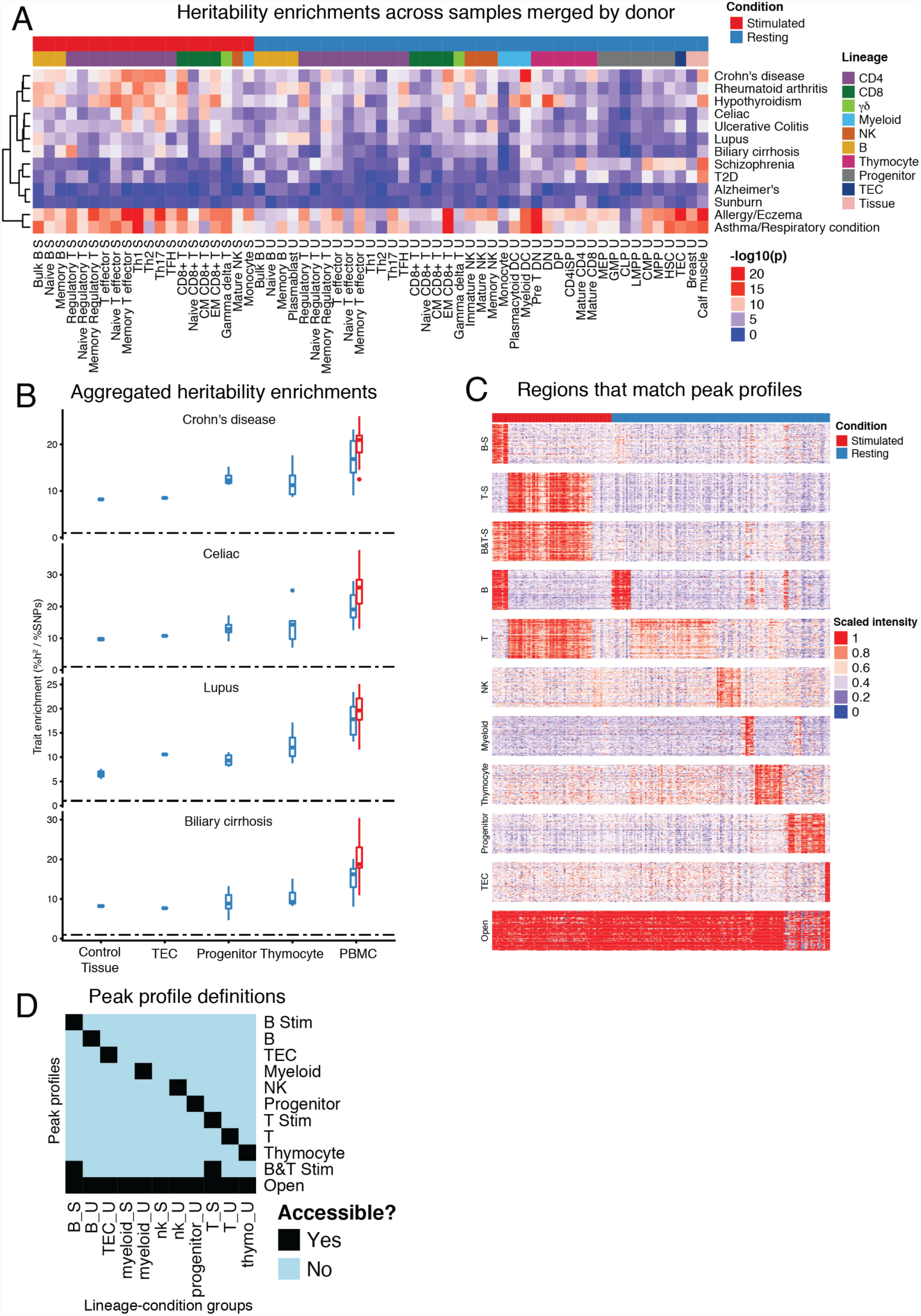
GWAS enrichment at chromatin accessible sites. (A) Significance of heritability enrichment between samples and complex traits. (B) Trait heritability enrichment of samples from various lineage and condition classes. Resting samples indicated in blue and stimulated samples in red. Innate cells were excluded. (C) Heatmap showing top 100 accessible peaks (y-axis) for lineage clusters (labeled) that are unique for different cell lineages conditions across samples (x-axis). Increasingly accessible regions are colored red corresponding to a larger scaled intensity value (see Methods). (D) Definition of peak profiles. Peak profiles are labeled on the y-axis, which are defined by accessibility in various lineage-condition groups on the x-axis.

**Figure S6.**
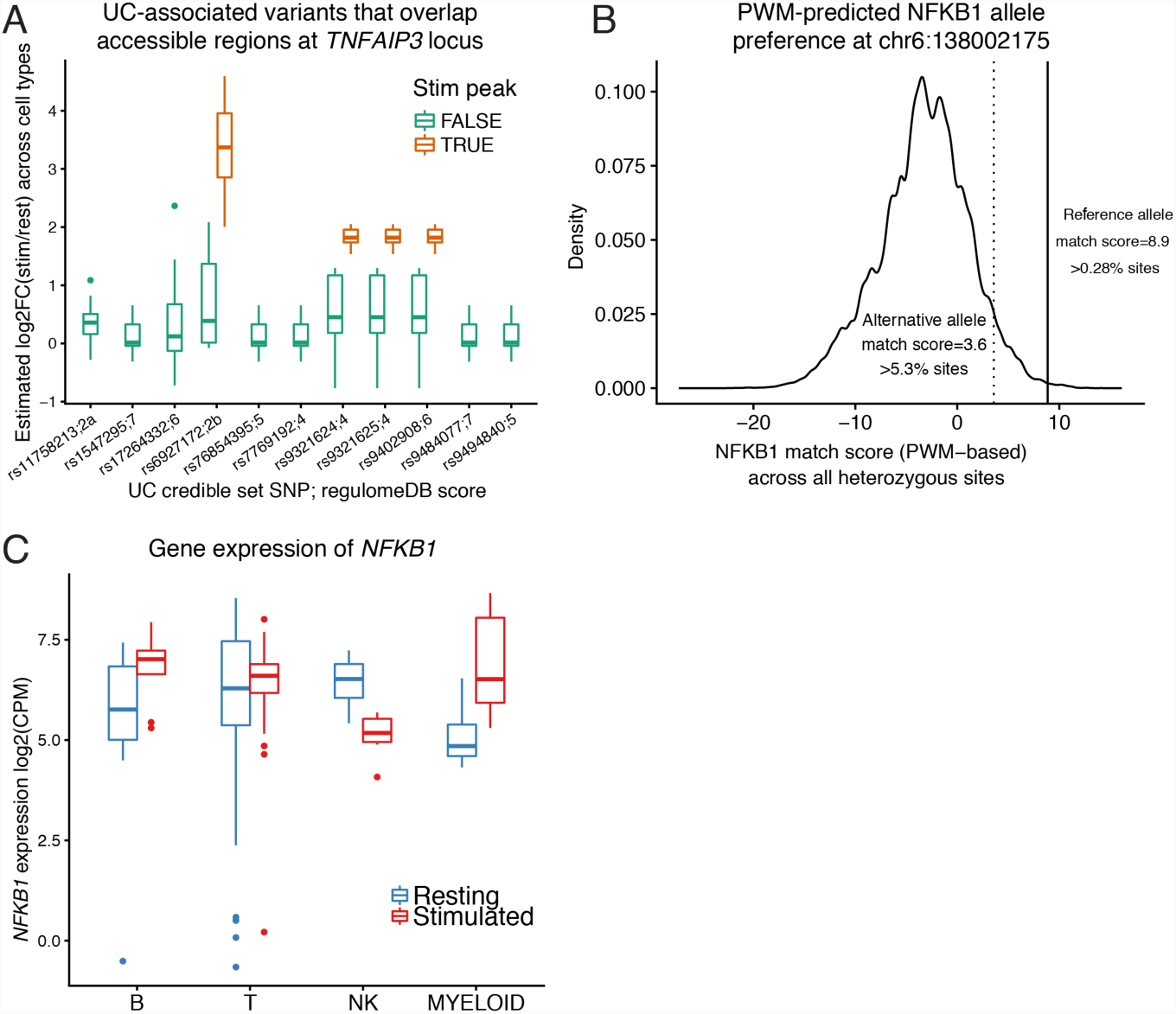
Putative regulation of TFNAIP3. (A) UC-associated variants that overlap accessible regions at the *TNFAIP3* locus and the distribution of stimulation-effects across all cell types. The variants rs9321624, rs9321625, and rs9402908 overlap the same accessible region. (B) The distribution of PWM-predicted NFKB1 binding affinity across all heterozygous sites from four donors. The reference allele at chr6:138002175 is represented by the solid vertical line, and the alternative is represented by the dotted vertical line. (C) Gene expression measured for *NFKB1* grouped by lineage and condition.

